# Paroxetine-induced transient apoptosis with delayed neurogenesis induces brain remodeling in developing zebrafish

**DOI:** 10.1101/2023.11.10.566506

**Authors:** Tomomi Sato, Kaito Saito, Tsubasa Oyu, Kazuna Yamaguchi, Maiko Osawa, Yuki Shiko, Yohei Kawasaki, Tomohiro Kurisaki, Sachiko Tsuda, Takeshi Kajihara, Masabumi Nagashima

## Abstract

Autism spectrum disorder (ASD) is a neurodevelopmental condition caused by various genetic and environmental factors. This disorder has the cardinal symptoms including impaired social behavior involving the amygdala. Ingesting antidepressants such as paroxetine in early pregnancy increases the risk of ASD in offspring. However, a comprehensive picture of the underlying pathogenic mechanisms remains elusive. Here we demonstrate that early exposure of zebrafish embryos to paroxetine retards neurogenesis in the optic tectum and the telencephalon that includes a region corresponding to the mammalian amygdala. Paroxetine-treated embryos exhibit impaired growth, with small heads and short body lengths resulting from transient apoptosis. This is reminiscent of the early-onset fetal growth restriction (FGR) associated with ASD. Interestingly, the delayed neurogenesis in the small heads was found to be restored after the cessation of paroxetine. This restoration was accompanied by extended retinotectal projections, suggesting brain-preferential remodeling. Finally, social behavior of the paroxetine-treated fish was examined. Our findings imply a biological mechanism underlying early exposure to paroxetine, a risk factor associated with the pathogenesis of ASD.

## Introduction

Autism spectrum disorder (ASD) is a neurodevelopmental disorder diagnosed in childhood [1]. The cardinal symptoms include impaired social behavior, communication disorder, and stereotyped repetitive behaviors [2]. To date, a large number of genes have been identified as related to ASD. These are involved in synaptic functions, transcriptional regulation, and brain development [1,3]. Furthermore, de novo mutations, paternal age, maternal infection and inflammation during pregnancy have also been implicated as causative factors [3–5]. Thus, various genetic and environmental factors contribute to the impaired brain development and functions observed in ASD [4–6]. Maternal depression and anxiety during pregnancy are also associated with an increased risk of ASD in offspring [7]. Selective-serotonin reuptake inhibitors (SSRIs), one of the conventional types of antidepressants, are prescribed to pregnant women based on risk-benefit considerations [8]. The drugs act on the maternal brain to exert their therapeutic effects but also pass through the placenta to the fetus [9,10]. Several studies suggest that maternal use of SSRIs during the first trimester of pregnancy increases the risk of ASD [11–14]. However, a comprehensive picture of the underlying pathogenic mechanisms remains elusive.

Fetal growth restriction (FGR) is a common complication during pregnancy and possibly caused by various maternal, placental, and fetal factors [15]. Early-onset FGR manifests as a morphologically symmetrical fetus with a small head and body due to fetal growth failure [16]. The causative factors include genetic abnormalities, congenital malformation, intrauterine infection, and teratogenic effects [15]. This type of FGR is also associated with ASD [17–19]. However, the pathogenesis of ASD and the pathophysiological role of SSRIs in the early-onset FGR is poorly understood.

Serotonin (5-hydroxytryptamine [5-HT]) is a monoamine neurotransmitter involved in emotional disorders such as depression and anxiety by modulating synaptic transmission in the brain [20,21]. SSRIs exert their therapeutic effects by inhibiting the reuptake of 5-HT via serotonin transporter (SERT), to sustain the concentration of extracellular 5-HT [22,23]. 5-HT also plays an important role in adult neurogenesis by promoting the self-renewal of neural stem cells in the hippocampus [24]. Moreover, maternal 5-HT synthesized in the placenta of pregnant mice has been found to transiently localize in the forebrain of early-stage embryos [25], raising the possibility that placenta-derived 5-HT regulates early brain development [26–28]. However, the effects of early-stage SSRI and 5-HT on embryonic brain development and postnatal behaviors are unclear. Elucidation of the primary causes is essential for early prediction and prevention of the risk for ASD. While SSRI during early pregnancy increases the risk of ASD, the primary cause remains ambiguous [11,13,14]. The involvement of various genetic and environmental factors makes it difficult to comprehensively understand the causes and consequences for the pathogenesis of ASD [1–4]. Thus, there is a need to elucidate the primary actions of SSRI on early brain development and the subsequent effects on social behavior. To address this question, we use zebrafish as a model of human embryos, because easy access to early embryos in a living and intact condition enables us to investigate the direct actions on those embryos. In zebrafish, various models for neurodevelopmental disorders have already been established, suggesting evolutionary conserved pathophysiological mechanisms between humans and zebrafish in the disorders [29–31]. We focus on the SSRI, paroxetine, because of its broad applicability across affective disorders, including major depressive and anxiety disorders [32]. In this study, we assess the effects of paroxetine and compare those with the control and 5-HT, on morphology, neurogenesis, projection in the brain during embryonic, larval stages, and finally on social behavior in juvenile stages.

## Results

### Amino acids interacting with paroxetine are completely conserved between human and zebrafish SERT

To investigate the effects of paroxetine on brain development in zebrafish embryos, we first examined whether the amino acid sequence of the serotonin transporter (SERT) targeted by paroxetine, is conserved between humans and zebrafish (Fig 1). Three-dimensional structures of paroxetine-binding sites and interacting amino acids in human SERT have already been well-defined by X-ray crystallography (Fig 1A) [33,34]. The entire amino acid sequence was 70% identical between the two species (Fig 1B). All the amino acids that interact with paroxetine at the central site of human SERT were found to be identical between the two species (Fig 1A, B). We then examined the expression of zebrafish SERT proteins by immunohistochemistry (Fig 1C and S1 Fig). SERT proteins were weakly but broadly expressed in the brains at 30 hours post-fertilization (hpf), at the time when pet1-positive serotonergic neurons develop in the raphe nuclei (S1 Fig) [35]. We found that SERT proteins were localized to the apical surfaces of radial glial cells in the ventricular zone at 24 hpf, implying that SERT play some roles in neural stem/progenitor cells.

**Fig 1.**
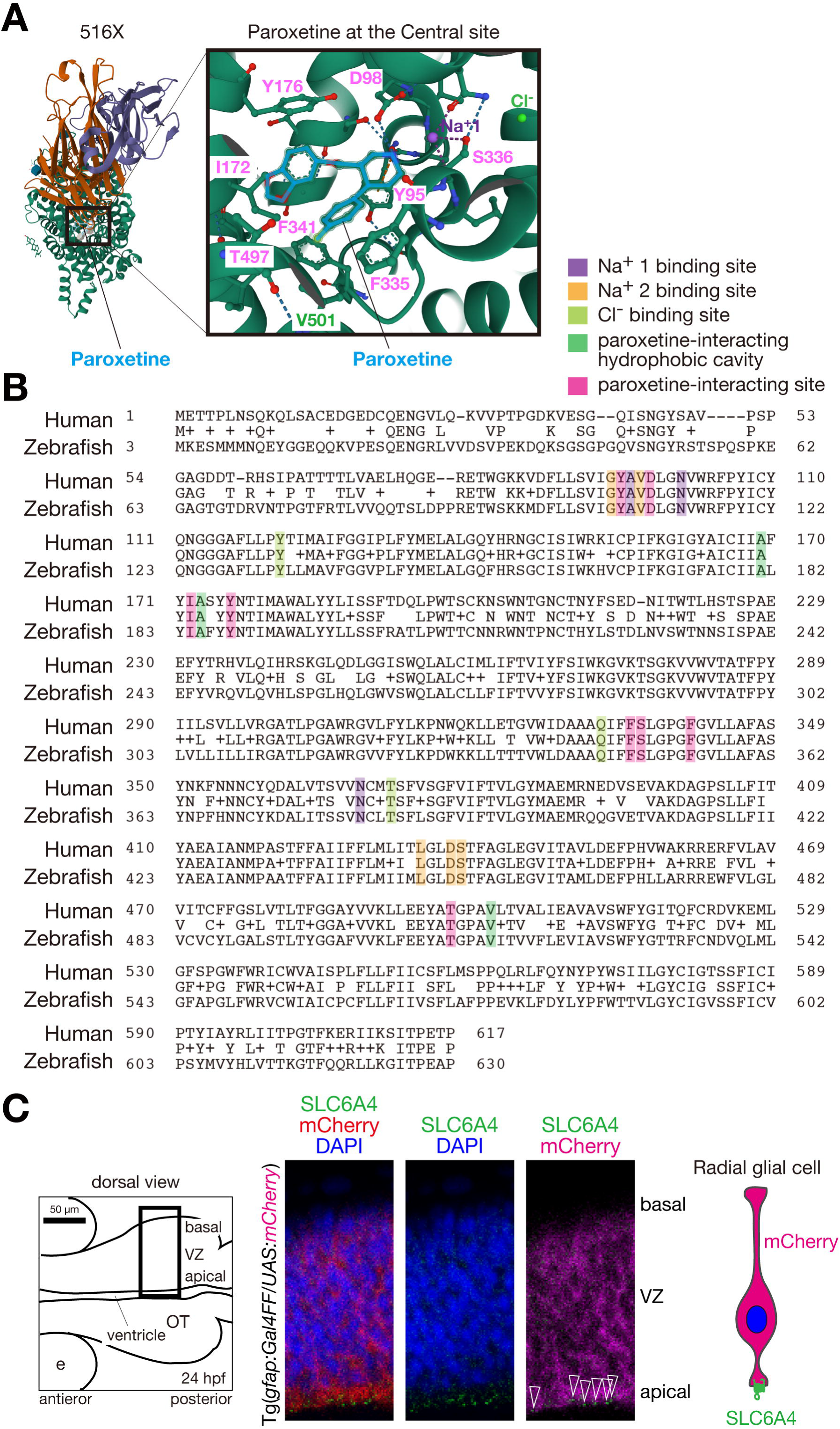
Amino acids interacting with paroxetine are conserved between human and zebrafish SERT. (A) X-ray structure of the ts3 human SERT complexed with paroxetine at the central site (516X, RCSB Protein Data Bank). Amino acids interacting with paroxetine (light blue), indicated in pink. An amino acid forming hydrophobic cavity, indicated in green. (B) Alignments of amino acid sequences of human and zebrafish SERT (P31645 and NP001035061). Amino acids interacting with paroxetine in the central site are completely identical between human and zebrafish. (C) SLC6A4 (SERT) proteins localized in the apical surface of the ventricular zone of the optic tectum (green, open arrowheads). A scheme of the embryonic brain in a dorsal view (Left). The corresponding region shown in the right at high magnification is indicated (Bold rectangle). A scheme of an RFP-expressing radial glial cell and the SLC6A4 localization is shown (Right).

### Early transient paroxetine generates embryos with a short body and a small head which is restored at later larval stages

To clarify the effects of paroxetine on embryonic development, we treated embryos with paroxetine and the effector 5-HT between 26–50 hpf, which roughly corresponds to the first trimester of human embryos (Fig 2A) [36]. To determine subthreshold concentrations appropriate for treatment, we treated embryos and calculated survival rates at different concentrations of paroxetine and 5-HT. Both compounds had no effect on the survival rate at less than 20 µM (Fig 2B). Then, we measured the head-tail body length and head area of paroxetine-treated embryos to parallel the indication of human fetal development estimated by crown-lump length (CRL) between weeks 8–11 of the first trimester in human pregnancy [37]. At first, we monitored the typical development of wild-type embryos. In these experiments, body lengths and head areas were gradually but significantly greater than those embryos at 50 hpf according to developmental stages (S2 Fig). We found that embryos treated with 10 µM paroxetine exhibited restricted growth with a slight but significant decrease in body length and head area at 50 hpf, whereas no significant decreases were detected in embryos treated with 1 µM paroxetine (Fig 2C-E). This closely resembled early-onset FGR of human embryos in the first trimester [16]. In contrast, 5-HT did not affect both the body length and head area of the embryos at the concentrations of 10 µM and 1 µM, indicating that the observed embryonic growth restriction is a paroxetine-selective action (Fig 2C-E). Embryos treated with 1 µM paroxetine for two days exhibited significantly shorter body lengths and smaller head areas at 73 hpf (S3 Fig). In addition, treatment with 10 µM paroxetine for one day had a similar effect on larval morphology at 3 dpf, suggesting that transient early-stage treatment is sufficient for paroxetine to exert its growth-inhibiting effects (S3 Fig). Therefore, we investigated the effects of early transient paroxetine on later development. 5-dpf larvae treated with 10 µM paroxetine during 26–50 hpf showed significant decreases in body lengths and head areas compared to the controls (S4 Fig). Interestingly, the paroxetine-treated larvae exhibited restoration of their head areas compared to the controls at 7 dpf despite persistent decreases in their body lengths (Fig 2F-I). This observation implied that brain-preferential remodeling occurrs between 5–7 dpf.

**Fig 2.**
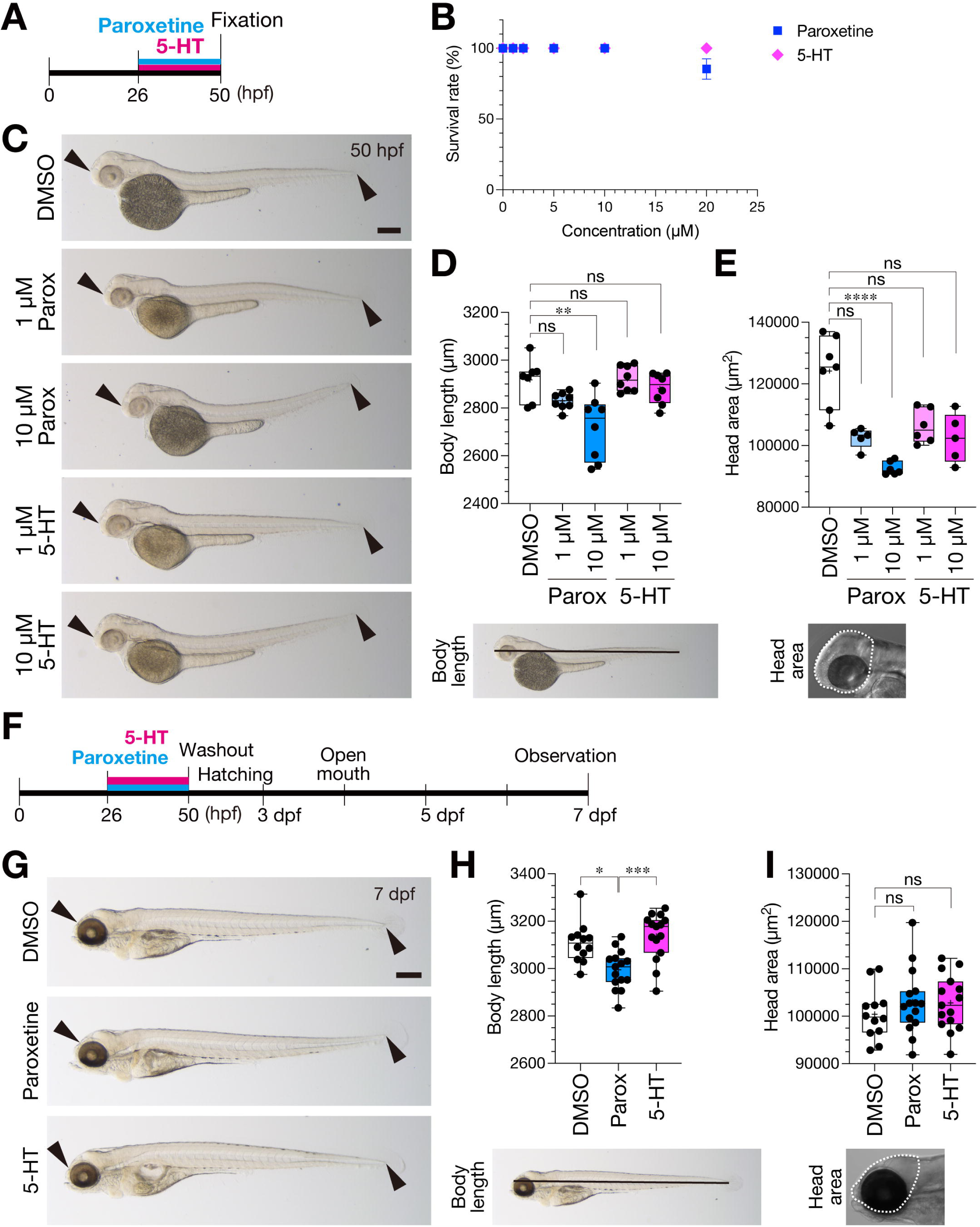
Early paroxetine induces embryonic growth restriction followed by preferential restoration in the head at a larval stage. (A) A scheme for the time course of treatment. (B) Dose-responses of survival rates for paroxetine- (blue) and 5-HT- (magenta) treated embryos. (C) Lateral views of 50-hpf zebrafish embryos treated with parox, 5-HT, or the control DMSO. (D, E) Quantification of the body lengths (D) and head areas (E) of the treated embryos. A representative image of an embryo indicating the quantified body length (black line, D) and head area (dotted line, E) are shown in the bottom. (F) Scheme for the treatment schedule and major events in zebrafish development. (G) Lateral views of 7-dpf larvae treated with parox, 5-HT, or DMSO. (H, I) Quantification of the body lengths (H) and head areas (I) of the treated embryos. A representative image of a larva indicating the quantified body length (black line, H) and head area (dotted line, I) in the bottom.

### Paroxetine but not 5-HT transiently induces local apoptosis in the head and tail tip

To elucidate the mechanisms underlying the different effects of paroxetine and 5-HT on embryonic growth and development, we examined the induction of apoptosis at 31 hpf, 5 hours after treatment with paroxetine or 5-HT by immunostaining with an antibody against active Caspase-3 (Fig 3A, B). We found a significant increase in the immunofluorescent signals of active Caspase-3 in the whole brain of paroxetine-treated embryos compared to the control dimethyl sulfoxide (DMSO)-treated embryos (Fig 3B, D, parox). The apoptosis induced by paroxetine was increased in a dose-dependent manner (Fig 3E). In contrast, no such increase was detected in 5-HT-treated embryos (Fig 3B, D, 5-HT). This is consistent with our observations that 5-HT had no effects on the head area (Fig 2C, E). Co-treatment of paroxetine with 5-HT also increased apoptosis in the brain, suggesting that the apoptosis is directly induced by paroxetine blocking the incorporation of 5-HT via SERT, but not through extracellular 5-HT (Fig 3F). Moreover, in addition to the apoptosis in the whole brain, apoptosis was also induced locally in the tail tip of paroxetine-treated embryos in an almost all-or-nothing manner (Fig 3C, G). Consistent with paroxetine-treated embryos, *slc6a4a* (*serta*)-knockdown embryos also exhibited growth restriction in the body length and increased apoptosis in the brain were observed in (S1 Fig). These results indicate that the increased apoptosis was caused by reduced 5-HT transportation due to the inhibition or knockdown of SERT. Thus. the local apoptosis in the brain and the tail tip corresponded to the growth restriction of the paroxetine-treated embryos with decreased head area and body length (Fig 2C-E). Next, we examined whether apoptosis persistently occurred in the paroxetine-treated brain at 48 hpf. A brain-wide decrease of Caspase-3-positive signals were detected in the paroxetine-treated embryos. Indeed, as representative brain regions, the signals were significantly decreased in the optic tecta (OT) and the telencephalons (T) of paroxetine-treated embryos, indicating that apoptosis acutely induced after the treatment was suppressed in the brains at later stages (Fig 3H-J).

**Fig 3.**
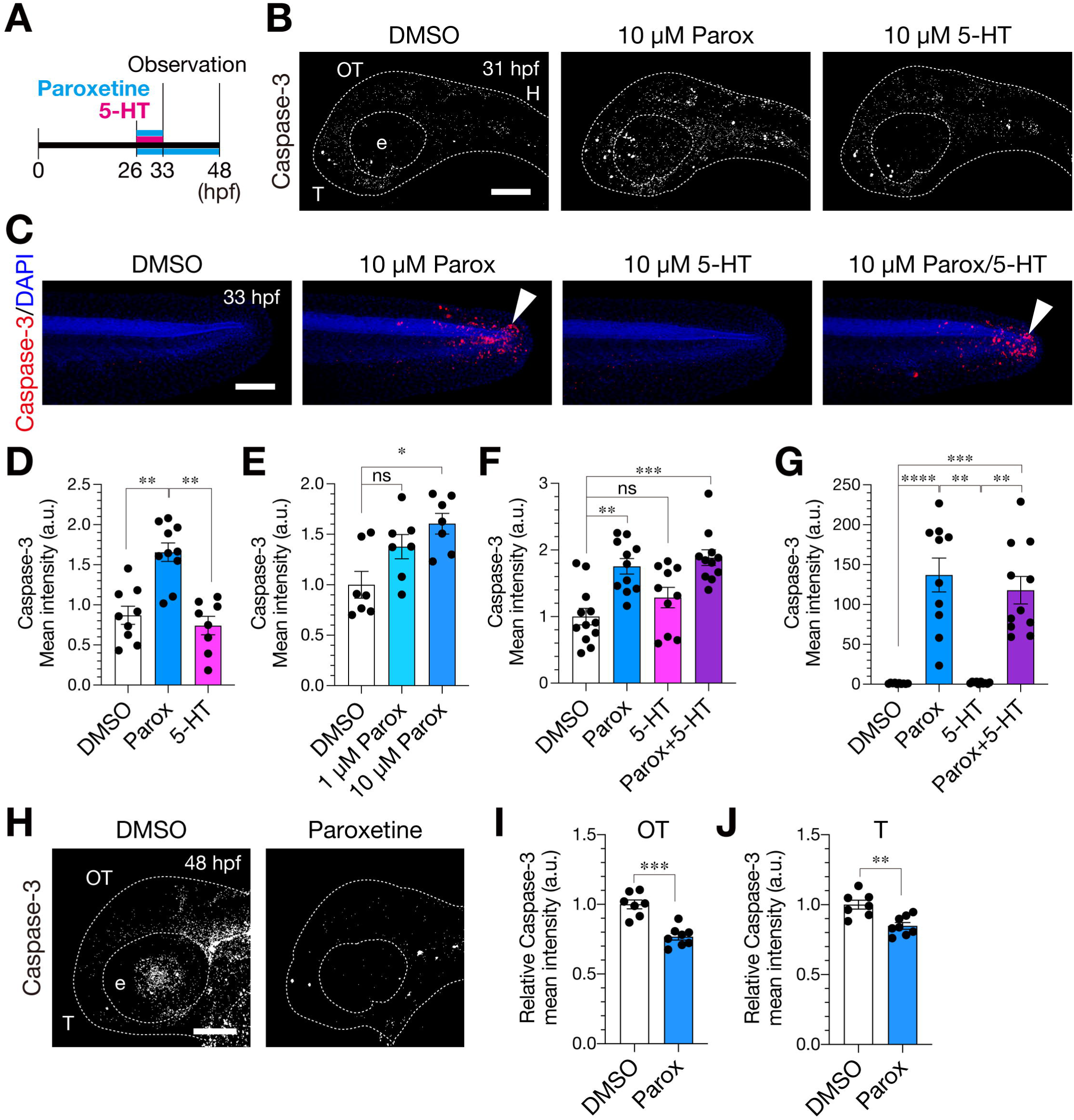
Paroxetine transiently induces local apoptosis in the brain and tail tip of treated embryos. (A) A scheme for the time course of treatment. (B) Lateral views of the heads of 31-hpf embryos treated with DMSO, parox, or 5-HT and immunostained for active Caspase-3 (white puncta). Images binarized above a threshold. Dotted line: Morphological outlines. (C) Lateral views of the tail tips of 33-hpf embryos treated with DMSO, parox, or 5-HT and immunostained for active Caspase-3. Anterior, left; posterior, right. (D-G) Quantification of the mean intensities of active Caspase-3 signals in the heads (D-F) and tail tips (G). (H) Lateral views of DMSO- or parox-treated embryos immunostained for active Caspase-3 (white puncta) at 48 hpf. Images binarized above a threshold. Dotted line: Morphological outlines. (I, J) Quantification of the mean intensities of active Caspase-3 in the optic tecta (I) and telencephalons (J) of the treated embryos. Scale bars: 100 µm (B, C, H). Data are presented as mean ± SEM. \**p* < 0.05, \*\**p* < 0.01, \*\*\*\**p* < 0.0001; ns, not significant. Kruskal-Wallis with Dunn’s multiple comparisons tests; parox, paroxetine; 5-HT, serotonin; DMSO, dimethylsulfoxide; hpf, hours post-fertilization; parox, paroxetine; SEM, standard error of the mean. OT, optic tectum; T, telencephalon.

### Paroxetine decreases post-mitotic neurons, whilst increasing proliferation of neural progenitor cells in the optic tectum

We then explored the effect of paroxetine on neurogenesis in the optic tectum, since it has been well defined as a visuomotor center in the zebrafish brain in terms of the development and function of the neural circuits [31, 38–42]. For this purpose, we used transgenic lines TgBAC(*neurod1:EGFP*), expressing enhanced green fluorescent protein (EGFP) in differentiated neurons [43], and Tg(*elavl3:kaede*), expressing Kaede under a pan-neuronal promoter *elavl3* (*huC*) [44] (Fig 4A and S4 Fig). To evaluate the effect on neurogenesis by treatment with paroxetine and other SSRIs using intact whole embryos, we examined fluorescence-positive area and the fluorescence intensity to establish simple indicators of the number of post-mitotic neurons expressing a fluorescent protein such as EGFP or Kaede. Previously we have revealed that GFP intensity in post-mitotic neurons increases with developmental stages, because it is proportional to the number and the time after differentiation of those neurons [31]. Imaging and relative quantification processes are shown in S4 Fig and Materials and Methods. We found that the EGFP and Kaede fluorescence in post-mitotic neurons were significantly decreased in the optic tectum and the telencephalon by the treatment with paroxetine during 26–50 hpf compared to the control (Fig 4A-C and S4 Fig). Indeed, the number of EGFP-positive neurons (with intensity above 70) in the optic tectum was significantly decreased in paroxetine-treated embryos at 50 hpf (Fig 4D), suggesting that the number of post-mitotic neurons was significantly decreased due to increased apoptosis by paroxetine (Fig 3B, D-F). Treatment with other SSRIs, fluoxetine, fluvoxamine and sertraline also showed decreased post-mitotic neurons in TgBAC(*neurod1:EGFP*) similar to that with paroxetine, indicating that the effects on post-mitotic neurons were shared with other SSRIs (Fig 4G and data not shown). Immunohistochemistry for CyclinD1 showed a significant decrease in the protein expression in the optic tectum of the SSRI-treated embryos (Fig 4H and S4 Fig). We then investigated the effect of upregulation of extracellular 5-HT by treating embryos with 5-HT during the same developmental stages. Similar to the paroxetine-treated embryos, EGFP fluorescence was significantly decreased in the optic tectum of 5-HT-treated embryos compared to the controls (Fig 4E, F), indicating that the effect of paroxetine on the post-mitotic neurons is recapitulated by 5-HT even in the absence of induction of apoptosis (Fig 3B, D, F).

**Fig 4.**
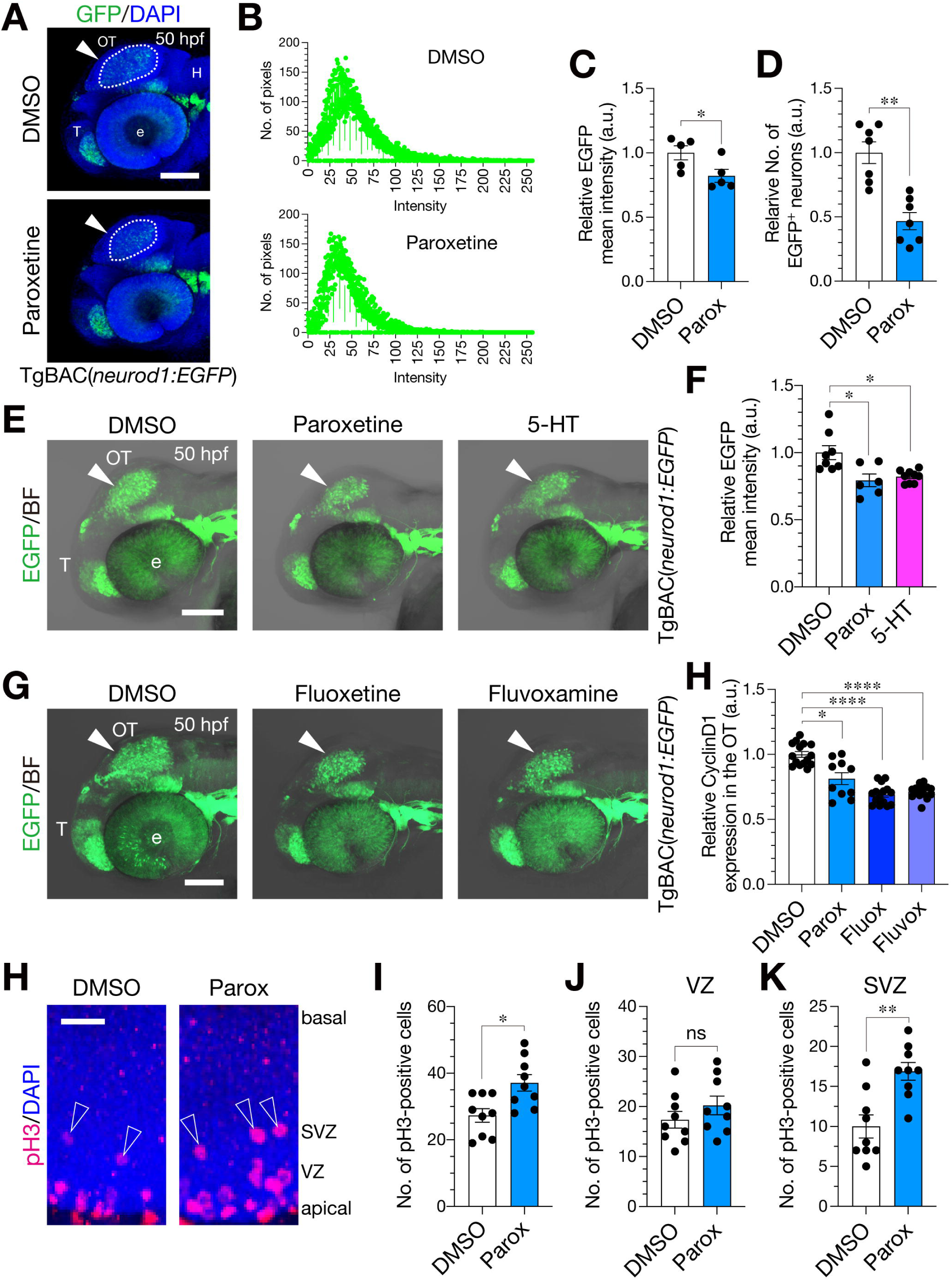
Paroxetine decreases post-mitotic neurons and promotes proliferation of neural progenitor cells in the optic tectum. (A) Representative 50-hpf TgBAC(*neurod1:EGFP*) embryos treated with parox and the control DMSO shown in lateral views. EGFP-positive region in the optic tectum (dotted ellipse) for quantification is indicated (arrowhead). (B) Intensity frequency of the EGFP-positive region in the optic tectum of DMSO- (top) and paroxetine- (bottom) treated embryos (DMSO, 24.38 ± 1.359 pixels; Parox, 21.30 ± 0.882 pixels; each n = 5). (C) Quantification of the mean EGFP intensity in the optic tectum (dotted ellipse with arrowheads in A). Unpaired t test. (D) Quantification of the number of EGFP-positive neurons with intensity above 70 in the optic tectum. (E) Lateral views of 50-hpf TgBAC(*neurod1:EGFP*) embryos treated with parox, 5-HT and the control DMSO. (F) Quantification of the mean EGFP intensity in the optic tecta (arrowheads in E) of the embryos. Kruskal-Wallis with Dunn’s multiple comparisons tests. (G) Lateral views of 50-hpf TgBAC(*neurod1:EGFP*) embryos treated with SSRIs (fluoxetine, fluvoxamine) and the control. (H) Quantification of CyclinD1 protein expression in the optic tectum treated with paroxetine, fluoxetine, fluvoxamine and the control. (I) High-magnification images of the optic tecta of zebrafish embryos treated with DMSO or parox and immunostained for pH3 (red), with DAPI (blue) co-staining for the nuclei. Open arrowheads, pH3-positive cells in the SVZ. (J-L) Quantification of the number of pH3-positive cells in the optic tecta (J), the ventricular zones (K), and the subventricular zones (L). Scale bars: 100 µm (A, E, G); 25 µm (I). Data are presented as mean ± SEM. \**p* < 0.05, \*\**p* < 0.01, \*\*\**p* < 0.001; \*\*\*\**p* < 0.0001; ns, not significant. Mann-Whitney U test; 5-HT, serotonin; DAPI, 4′,6-diamidino-2-phenylindole fluorescent stain; DMSO, dimethylsulfoxide; e, eye; EGFP, enhanced green fluorescent protein; H, hindbrain; hpf, hours post-fertilization; OT, optic tectum; parox, paroxetine; fluox, fluoxetine; fluvox, fluvoxamine; pH3, phospho-histone H3; SEM, standard error of the mean; SVZ, subventricular zone; T, telencephalon; VZ, ventricular zone.

In adult neurogenesis of both rodents and humans, SSRIs are known to promote the proliferation of neural stem cells [24,45]. Then, we examined the proliferation of neural stem/progenitor cells in the optic tectum by immunostaining with anti-phospho-histone H3 (pH3) antibody to label mitotic cells [31]. We found that paroxetine-treated embryos showed an increase in the number of pH3-positive mitotic cells at 50 hpf (Fig 4I, J). The increase was selectively observed in the subventricular zone (SVZ) where cell divisions of intermediate progenitor cells occur for neurogenesis and proliferation (Fig 4I, K, L) [46]. Expression of SOX2, a high-mobility group (HMG)-box transcription factor to suppress cell differentiation and promote self-renewal in proliferating neural stem/progenitor cells, was enhanced in paroxetine-treated embryonic brains, with a concomitant decrease in Kaede-expressing neurons at 48 hpf, although an increase in *sox2* mRNA expression was not detected by RT-qPCR with whole embryos (S4 Fig) [47]. These results suggest that paroxetine promotes the proliferation of neural progenitor cells in the optic tectum.

### The development of retinotectal projections is impaired by early transient paroxetine and later restored with extended retinal axons

To investigate whether the decreased neurons in the brain of paroxetine-treated embryos impacts later neural circuit formation, we examined the retinotectal projections. We used transgenic line Tg(*pou4f1-hsp70l:GFP*) expressing GFP in excitatory tectal neurons and retinal ganglion cells (RGCs), and line Tg(*pou4f3:GAL4;UAS:mCherry*) expressing mCherry in RGCs [43,44]. In wild-type larvae, GFP- and mCherry-expressing areas gradually and significantly increased with developmental stages in the optic tecta compared with those areas at 72 hpf (S2 Fig). In contrast, the fluorescence of both the GFP and mCherry-expressing areas was significantly decreased between 5 and 7 days post fertilization (dpf) during which activity-dependent competition followed by dynamic synapse formation or elimination occurs in retinal axons and tectal dendrites, respectively (S2 Fig) [38,39]. Treatment with 1 µM paroxetine for two days caused significant decreases in both the area and the fluorescence of the region expressing GFP and mCherry in the optic tectum at 73 hpf (Fig 5A-E). One-day treatment with 10 µM paroxetine also showed a similar effect on the GFP-expression region (S3 Fig). Thus, transient early-stage treatment of paroxetine is responsible for suppressing the development of the retinotectal projections, consistent with the growth-inhibiting effects on larval morphology (S3 Fig). Therefore, we examined the effects of the early transient paroxetine on later development of the retinotectal projections. In accordance with the restoration of the head area at 7 dpf (Fig 2G, I), paroxetine-treated 7-dpf larvae exhibited restoration of both the area and the fluorescence of the GFP-expressing region and the fluorescence of the mCherry-expressing region in the optic tectum by reverting the values to those in the controls (Fig 5F-H, J). Furthermore, the areas of mCherry-expressing regions were significantly expanded in paroxetine-treated larvae compared to the controls (Fig 5I, parox). This expansion was recapitulated in 5-HT-treated larvae (Fig 5I, 5-HT), suggesting that early transient paroxetine drives the extension of retinal axons through 5-HT at later neurodevelopmental stages.

**Fig 5.**
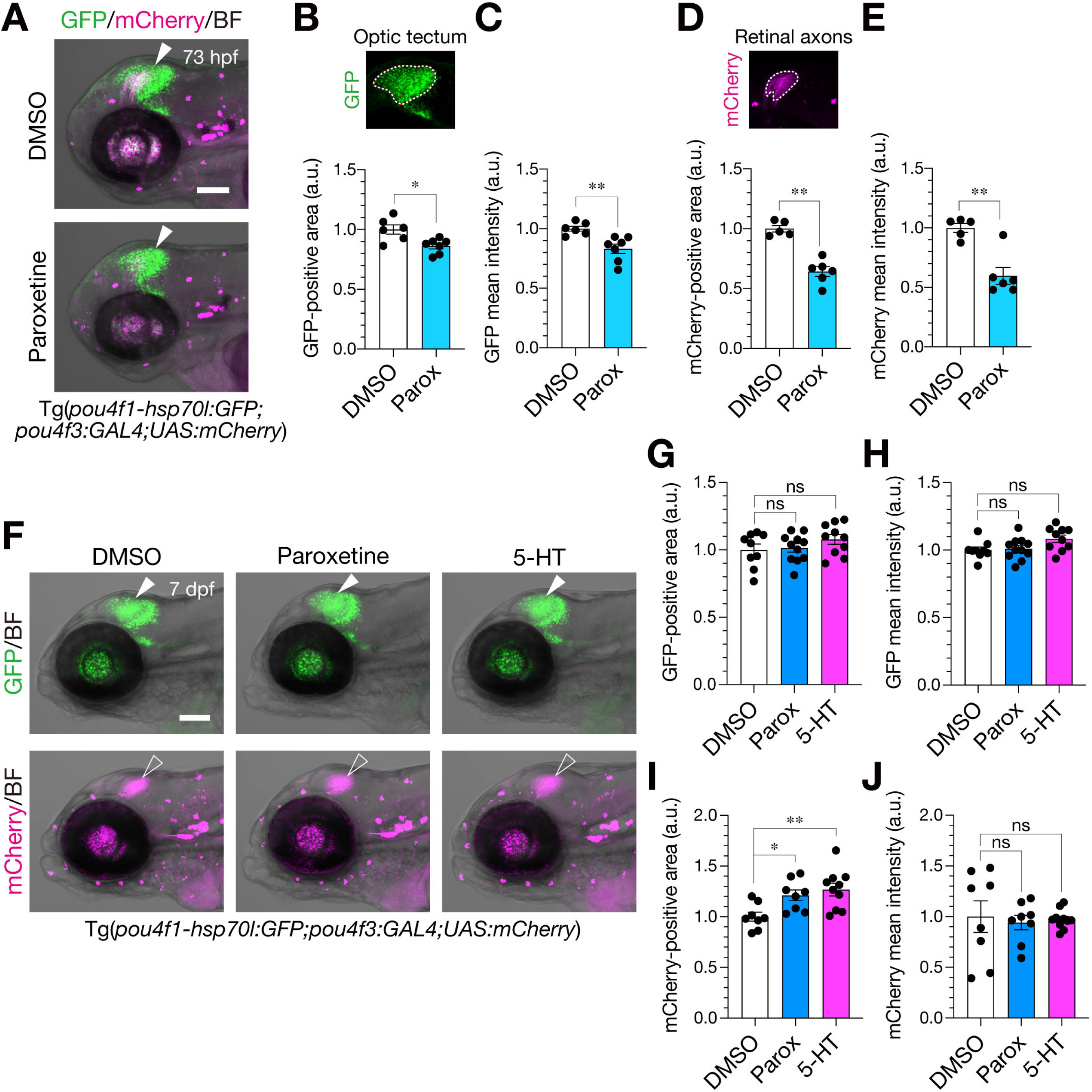
Early transient paroxetine inhibits the development of the retinotectal projection leading to brain-preferential remodeling. (A) Lateral views of the heads of 73-hpf Tg(*pou4fl-hsp70l:GFP; pou4f3:GAL4;UAS:mCherry*) embryos treated with DMSO or parox at 1 µM for 2 days from 26 hpf. Tectal neurons and retinal axons express GFP (green) and retinal axons express mCherry (magenta). (B) Quantification of the GFP-positive areas. (C) Mean GFP intensities in the optic tecta of the transgenic embryos. (D) Quantification of the mCherry-positive areas. (E) Mean mCherry intensities in the optic tecta of the transgenic embryos. A representative image indicating the quantified area (dotted line) in the optic tectum are shown (B, D, top insets). Mann-Whitney U test. (F) Lateral views of the heads of 7-dpf Tg(*pou4fl-hsp70l:GFP;pou4f3:GAL4;UAS:mCherry*) larvae treated with parox, 5-HT, or the control DMSO. Optic tecta (filled arrowheads) and retinal axons (open arrowheads). (G-J) Quantification of the GFP-positive areas (G), the mean GFP intensities (H), the mCherry-positive areas (I), and the mean mCherry intensities (J) in the optic tecta of the transgenic larvae. Kruskal-Wallis with Dunn’s multiple comparisons tests; Scale bar, 100 µm. Data are presented as mean ± SEM. \**p* < 0.05, \*\**p* < 0.01; ns, not significant. 5-HT, serotonin; DMSO, dimethylsulfoxide; GFP, green fluorescent protein; hpf, hours post-fertilization; parox, paroxetine; SEM, standard error of the mean.

### Early paroxetine negatively impacting neurogenesis in the telencephalon

Next, we investigated whether the early paroxetine treatment affects the development of the telencephalon that includes a region corresponding to the mammalian amygdala [29]. The amygdala plays a critical role in social behavior. Amygdala dysfunction causes impaired social behavior characteristic of ASD [2]. To assess the effects of paroxetine, SAGFF120A gene-trap line was used, because the line locally expresses Gal4FF in neurons of the telencephalon which are essential to active avoidance fear conditioning [49]. Fluorescence of the GFP-expressing areas was significantly decreased in the telencephalons of paroxetine- and 5-HT-treated embryos compared to the control at 76 hpf (parox, *p* = 0.0279; 5-HT, *p* = 0.0250, Fig 6B-E) as observed in the optic tecta, although the decrease was significant for 5-HT but not for paroxetine at 48 hpf (parox, *p* = 0.3344; 5-HT, *p* = 0.0006, S6 Fig). Consistently, the GFP fluorescence in the telencephalon was restored at 7 dpf following washout of paroxetine and 5-HT at 2 dpf (Parox, *p* = 0.1419; 5-HT, *p* = 0.6623; Fig 6C-F). This restoration concomitantly occurred with that in the head area, corresponding to the results observed in the larval optic tecta (Fig 2I, Fig 5F, H, J, S6 Fig).

**Fig 6.**
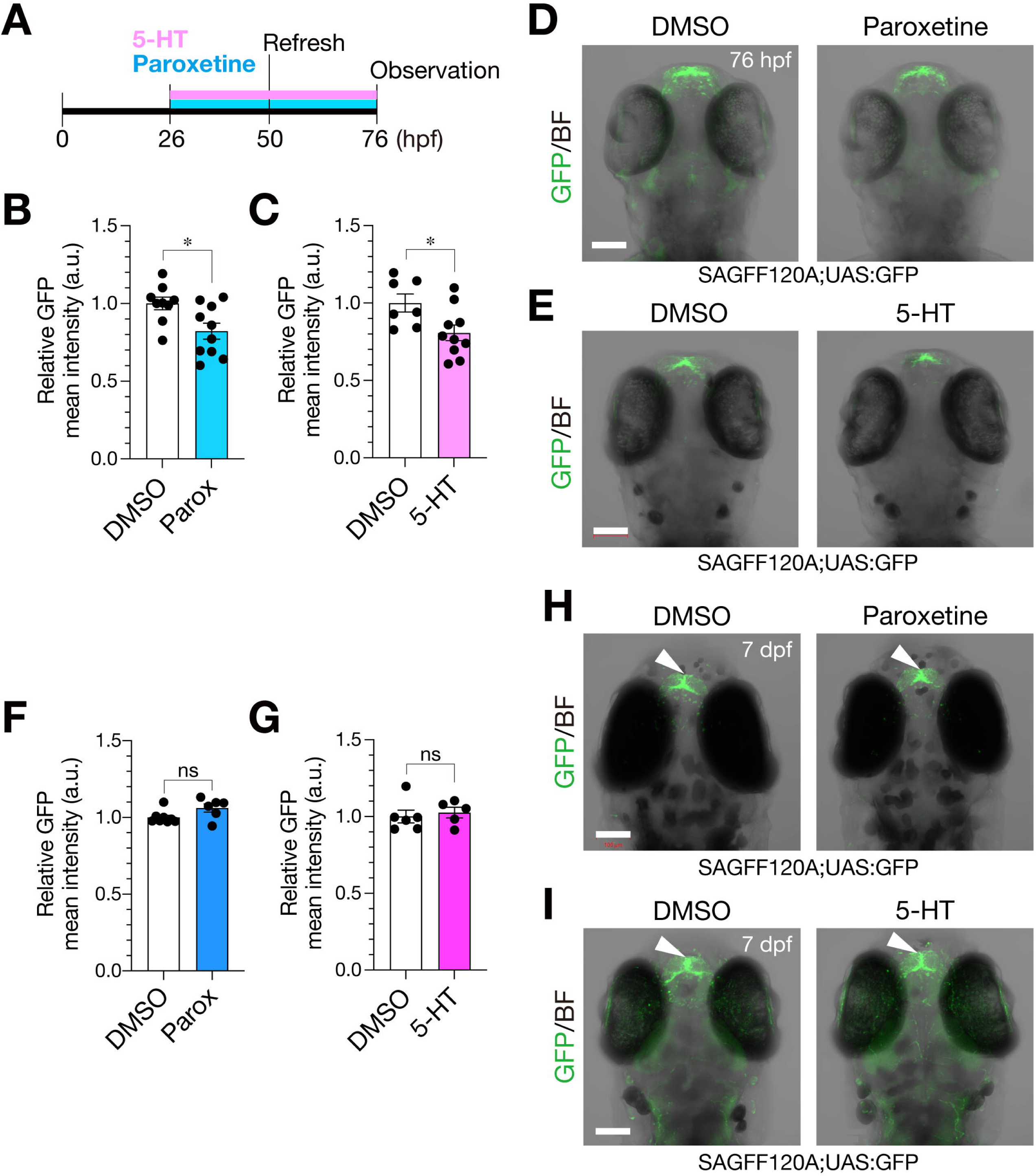
Early transient paroxetine exerts suppressive effects on neurogenesis followed by restoration in the telencephalon. (**A**) A schematic diagram of treatment with 1 µM paroxetine. (**B**, **C**) Quantification of GFP intensity in the telencephalon of the paroxetine-treated (**B**) and 5-HT-treated larvae (**C**). (**D**, **E**) Dorsal views of 76-hpf SAGFF120A;UAS:GFP larvae treated with paroxetine (**D**), 5-HT (**E**), and the control DMSO. (**F**, **G**) Quantification of the mean GFP intensities in the telencephalons of the paroxetine-treated (**F**) and 5-HT-treated larvae (**G**). *p* = 0.1419 (**F**), *p* = 0.6623 (**G**), Mann-Whitney U test. (**H**, **I**) Dorsal views of 7-dpf SAGFF120A;UAS:GFP larvae treated with paroxetine (**H**), 5-HT (**I**), and the control DMSO. Scale bar, 100 µm. Data are presented as mean ± SEM. \**p* < 0.05, \*\*\**p* < 0.001; ns, not significant. 5-HT, serotonin; DMSO, dimethylsulfoxide; GFP, green fluorescent protein; hpf, hours post-fertilization; parox, paroxetine; SEM, standard error of the mean.

Finally, we explored whether these neurodevelopmental effects of early transient paroxetine have an impact on juvenile social behavior. This is because no obvious differences in food intake and feeding behavior were observed between paroxetine-, 5-HT-treated and the control fish. Indeed, the body length of paroxetine-treated fish was not significantly different from that of the control fish (data not shown). To assess social behavior of a young fish, we developed a behavioral apparatus using a test arena separated into three chambers: a test area, a sibling area, and a non-sibling area (Fig 7A; see Methods). The swimming trajectories of the fish in the test area were recorded following habituation in the absence (non-sibling trial) and presence (sibling trial) of siblings in the sibling area (Fig 7B). We examined which area the fish was located in the sibling (blue) or non-sibling area (red) per frame to determine whether the fish show a social preference for the sibling frame. This was measured by calculating the social preference index (SPI, -1 < SPI < 1) (Fig 7C; S1-S6 Videos; see Methods) [50]. Total swimming distances showed no significant differences between paroxetine-, 5-HT-treated and the control DMSO-treated fishes both in non-sibling and sibling trials (Fig 7D). The mean SPI difference between the sibling and non-sibling trials tended to be more negative than the control for both paroxetine and 5-HT, although comparisons of paroxetine and 5-HT with the control were not significantly different (paroxetine, *p* = 0.103, 5-HT, *p* = 0.083, permutation test) (Fig 7E). The behavioral analysis showed that the incidence of positive SPI (sibling trial > non-sibling trial, white) was lower in the fish treated with paroxetine and 5-HT than in the control fish, whereas that of negative SPI (sibling trial < non-sibling trial, black) was increased in the paroxetine- and 5-HT-treated fish compared to the control (Fig 7F; DMSO, n = 13; parox, n = 7; 5-HT, n = 6). This was also not significantly different in comparison with the control (paroxetine, *p* = 0.195, 5-HT, *p* = 0.096, permutation test). However, these results raised the possibility of a tendency for negative social preference in those fishes. On the other hand, we found a difference in the effect on total swimming distance between paroxetine and 5-HT. The total swimming distances were significantly decreased in sibling trials of the control and paroxetine-treated fish, but not in 5-HT-treated fish (Fig 7G). Thus, the total swimming distances of the control exhibited negative values for the differences between sibling and non-sibling trials (Fig 7H). This was observed in paroxetine-treated fish, whereas the decrease in the distance was abolished and the difference was significantly increased in the 5-HT-treated fish compared to the control and paroxetine (control, *p* = 0.0427; parox, *p* = 0.0082; Fig 7G, H). These results indicated a difference in the effects on behavior by early transient exposure between paroxetine and 5-HT.

**Fig 7.**
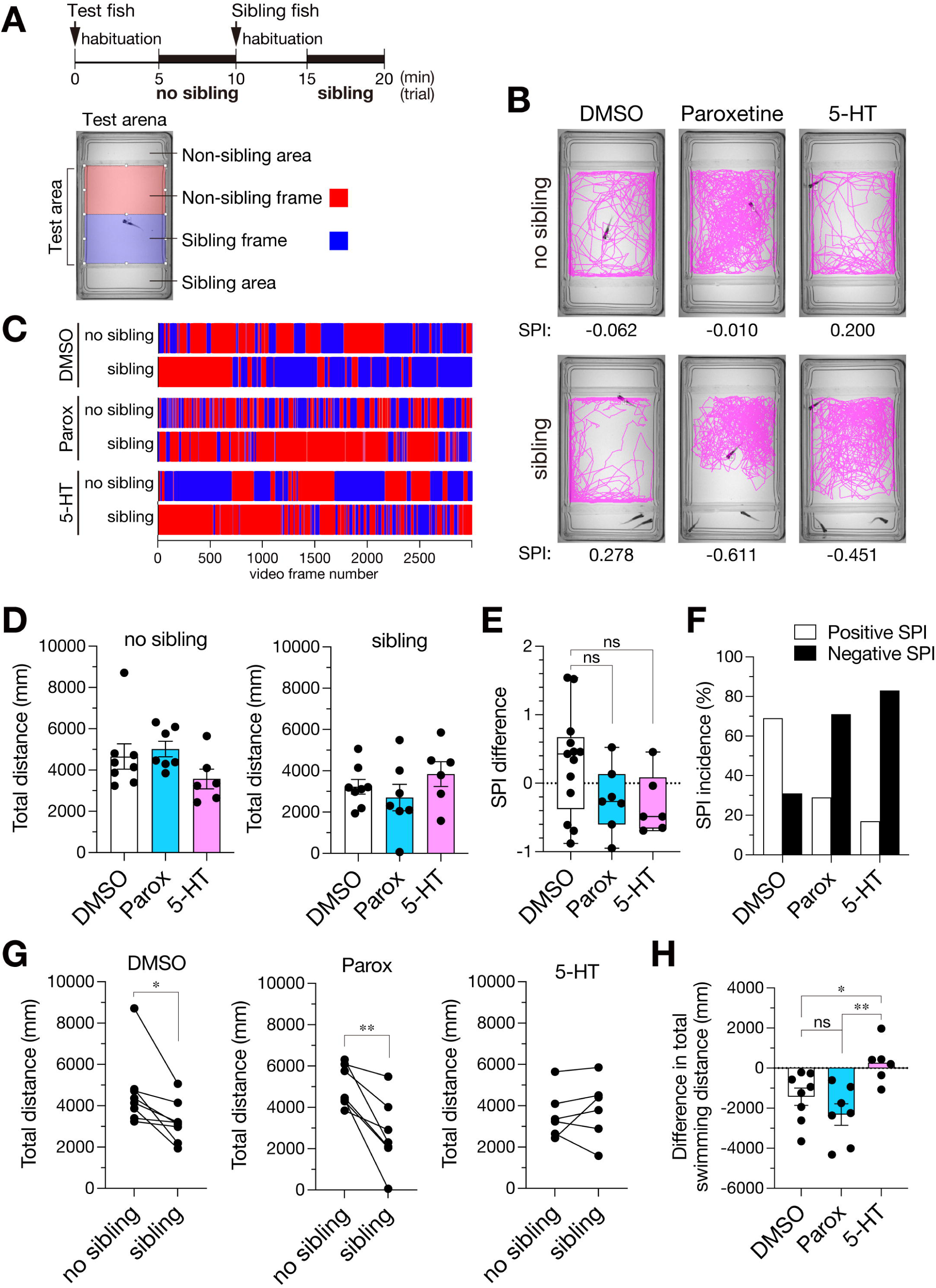
Social behaviors of early transient paroxetine- and 5-HT-treated zebrafish. (A) (top) A schematic diagram of the social behavioral test. (bottom) A test arena used in this study. The sibling frames and non-sibling frames are represented in blue and red, respectively. (B) Representative swimming trajectories of the fish treated with DMSO (control), parox, or 5-HT in the non-sibling (top) and sibling trials (bottom). The SPIs indicated below. (C) A total of 3,000 frames recorded over 5 min for the trials shown in (B) with color-coded according to the frames in (A). (D) Total swimming distances of DMSO-, Parox- and 5-HT-treated fish in non-sibling (left) and sibling (right) trials. (E) Differences in SPIs between the sibling and non-sibling trials. Data are presented as mean ± SEM, Permutation test. (F) Incidences of positive SPI (sibling > non-sibling, white) and negative SPI (sibling < non-sibling, black) in the fish treated with DMSO, parox, and 5-HT (n = 13, 7, 6, respectively). (G) Changes in the total swimming distances between non-sibling and sibling trials for DMSO-, Parox- and 5-HT-treatd fish. (H) Differences in the total swimming distances of the fish between the sibling and non-sibling trials. Data are presented as mean ± SEM, \**p* < 0.05, \*\**p* < 0.01; ns, not significant. Brown-Forsythe with Dunnett’s multiple comparisons tests. 5-HT, serotonin; ANOVA, analysis of variance; DMSO, dimethylsulfoxide; GFP, green fluorescent protein; hpf, hours post-fertilization; parox, paroxetine; SEM, standard error of the mean; SPI, social preference index.

## Discussion

In this study, we show that early exposure to paroxetine restricts embryonic growth, resulting in short body lengths and small heads in zebrafish. In humans, early-onset FGR manifests as symmetrical morphology with a small head and body, and the causative factors include teratogenic effects [15,16]. Thus, the paroxetine-treated zebrafish embryos are considered as a model of early-onset FGR in humans. The growth restriction is induced by paroxetine but not by 5-HT which is incorporated into cells through SERT, the target protein blocked by paroxetine. Consistently, apoptosis is locally induced by paroxetine in the brain and the tail tip, but not by 5-HT, suggesting that apoptosis is the underlying mechanism for the paroxetine-induced embryonic growth restriction. However, these results imply disparities between the effects of paroxetine and 5-HT on embryogenesis. SSRIs including paroxetine have already been reported to induce apoptosis in various malignant cell types, including gliomas, neuroblastomas, leukemia, lymphomas, and colon tumors, in addition to normal cells such as myeloid cells, osteoblasts, and osteoclasts [51–54]. These lines of evidence suggest a shared mechanism of action of SSRIs for the apoptosis induction in proliferating cells, potentially attributable to the etiology of adverse effects of SSRIs on embryogenesis. In addition, another SSRI, fluvoxamine, has recently been identified that have therapeutic potential as an immunomodulator due to its ability to suppress immune responses which is likely to reduce the risk of deterioration in COVID-19 [55,56]. These pleiotropic effects of SSRIs are reminiscent of those of thalidomide.

Thalidomide also has proapoptotic properties to exert the teratogenic effects on embryogenesis, and the exposure during early pregnancy has been associated with ASD in the offspring [57,58]. Furthermore, the thalidomide derivatives are therapeutically used as immunomodulatory drugs [59,60]. Taken together with those studies, our findings imply a shared common mechanism underlying teratogenic and therapeutic effects of SSRI which are related to the pathogenesis of ASD.

Here, we reveal that paroxetine- and 5-HT-treated embryos exhibit decreased post-mitotic neurons in the optic tectum. The effect of paroxetine is reproduced by the treatment with other SSRIs, suggesting that those effects are attributable to a shared property of antidepressants, serotonin re-uptake inhibition. The SSRI-treated embryos also show a significant decrease in CyclinD1 protein expression. These results imply that neurogenesis is suppressed by SSRIs, because Cyclin D1 play a role in promoting neurogenesis as a cell cycle-independent function [71]. Despite those effects on post-mitotic neurons, the number of pH3-positive mitotic cells are selectively increased in the subventricular zone. These observations raise the possibility that the proliferation of neural stem/progenitor cells is promoted by paroxetine and increased 5-HT at the expense of the generation of neurons from neural stem/progenitor cells during brain development [24,31,61]. This is supported by increased expression of SOX2, a transcription factor regulated by the PI3K/AKT signaling pathway that is not only inevitable for the induction and maintenance of stem cells, but also a marker of proliferating neural stem/progenitor cells in the brain [62,72]. Our observations are consistent with the effect of SSRI that increases the number of neural stem/progenitor cells in the dentate gyrus of the adult mammalian hippocampus through the increased concentration of 5-HT [24,45,63]. The effect of SSRI is thought to represent the underlying mechanism of the therapeutic effect on the maternal brain. On the other hand, our results suggest a biological mechanism that underlie an adverse effect on the embryonic brain which may be caused by SSRI ingestion during pregnancy [11–14] (S7 Fig). Indeed, we show that apoptosis is transiently induced and thereafter decreased in the brain, which generates growth restriction of paroxetine-treated embryos. These observations are consistent with a previous report showing that SERT-knockout mice exhibit reductions in programmed cell death in the brain [64]. As for the impact of the adverse effect, this apoptosis in the embryonic brain *per se* may elicit maternal inflammation through damage-associated molecular patterns (DAMPs) [73]. However, at present, it remains unclear how paroxetine induces apoptosis. Our results showing a similar effect of SSRIs on the post-mitotic neurons, that is decreased neurogenesis, suggest that incorporation of serotonin through SERT will be critical for survival of neural stem/progenitor cells leading to generation of neurons (S7 Fig).

Although there is a difference in apoptosis induction at early stages between paroxetine and 5-HT, both of which cause delayed neurogenesis. This implies that the delayed neurogenesis is a leading cause of the later effects on the retinotectal projection and social behavior, because both the treatments show a similar tendency for significant extension and negative SPI, respectively. Intriguingly, we find that the head size is restored at a later larval stage in early paroxetine-treated embryos, whereas body length is not, implying brain-preferential remodeling. Likewise, our data support that the delayed neurogenesis in the optic tectum decreases the areas and the fluorescence in the retinotectal projection of the paroxetine-treated embryos, which are later restored after the removal of paroxetine. These observations raise the possibility that remodeling occurs in neural networks in the larval brains during the restoration processes, because the remodeling period corresponds to the activity-dependent refinement stages in the retinotectal projection [38,39]. Indeed, we reveal extended projections of retinal axons in the restored brain. This phenotype in axonal projections is similar to that observed in SERT-knockout mice which exhibit impaired segregation of retinal axons both in the superior colliculus and the dorsal lateral geniculate nucleus, leading to the disruption of sensory map formation [65,66]. Those extended axons are also observed in the optic tecum treated with 5-HT. Similarly, the decreased neurogenesis is detected in both the optic tecta treated with paroxetine and 5-HT. Thus, our findings suggest that decreased neurogenesis is a cause of extended axonal projections, because those projections are observed regardless of whether growth restriction and local apoptosis are induced by the treatments. This provides insight into a biological basis for the neurodevelopmental disorders associated with early-onset FGR. The decreased neurogenesis followed by the restoration is also observed in the telencephalon treated with paroxetine and 5-HT using the gene-trap line SAGFF120A. The extent of the decrease in the fluorescent protein expression is different in the telencephalon of SAGFF120A from that of Tg(*elavl3:Kaede*) at 48 hpf. This is likely due to the difference in the population of neurons labeled by the fluorescent proteins. In the gene-trap line, neurons in the telencephalon that includes a region corresponding to the mammalian amygdala are visualized by GFP. These neurons have been identified that play a critical role in active avoidance fear conditioning [29,49]. Therefore, the restoration in the telencephalon implies that the treatments cause remodeling similarly to the optic tectum, which affects the neural circuit function involved in fear conditioning. Indeed, in individuals with ASD, the number of neurons in the amygdala has been found not to undergo neurotypical alterations during their development [67].

Finally, we investigate the impacts of delayed neurogenesis on the development of social behavior. In the control fish, the SPI differences between sibling and non-sibling trials vary among individuals, and thereby these are no significant difference in paroxetine- and 5-HT-treated fish as compared with the control. However, the incidence of negative SPI (sibling > non-sibling trials) is markedly increased in both the treated fish. No significant differences in the total swimming distances between paroxetine-, 5-HT-treated and the control fishes would exclude the possibility that the changes in social behavior are due to those in locomotor activity. A significant decrease in sibling trials of the control and paroxetine-treated fish can be explained by habituation, but not explain a similar tendency for increased negative SPI in Paroxetine- and 5-HT-treated fish. This will exclude the possibility of habituation. In addition, our observations of no obvious defects of the treated fish in food intake and feeding behavior which mainly depend on the visual system mostly rule out the possibility of visual impairment [74]. Therefore, our observations raise the possibility of a tendency to negatively impact the development of social behavior by early exposure to paroxetine and 5-HT. However, these early environmental conditions per se are less likely to be sufficient to have a significant effect on social behavior, because the effects on SPIs do not show significant differences. Nevertheless, transient apoptosis with delayed neurogenesis can be at risk for neurodevelopmental disorders, even when that is induced by other genetic or environmental factors. Of note, the effects on total swimming distance are significantly different between paroxetine and 5-HT, implying a difference in the effects on neural circuit functions. These findings may offer a biological mechanism underlying early risk factors associated with the pathogenesis of ASD. However, a more detailed analysis is a prerequisite for fully understanding the difference in the effects of paroxetine and 5-HT on the structure and function of the relevant neural circuits. In conclusion, our study provides evidence for the effects of early transient paroxetine and 5-HT on embryonic brain development and behaviors. Furthermore, this will represent a potential simple screening system for evaluating risk factors for ASD and neurodevelopmental disorders affecting social behavior using zebrafish.

## Methods

### Zebrafish maintenance and strains

Wild-type and transgenic zebrafish were kept at 28°C under 14 hours:10 hours light/dark cycles as described previously [68], in an IWAKI Small fish breeding unit LAbREED ITS. To obtain transgenic embryos, a wild-type strain Riken WT (RW) was crossed with transgenic lines as follows: Tg(*elavl3:Kaede*) to visualize post-mitotic neurons [44]; TgBAC(*neurod1:EGFP*) to visualize differentiated neurons[43]; Tg(*pau4f1-hsp70l:gfp*) to visualize development of post-mitotic excitatory neurons and retinal axons in the optic tectum [40]; Tg(*pau4f3:GAL4;UAS:mCherry*) to visualize retinal axons in the optic tectum [48]; and SAGFF120A;UAS:GFP to visualize neurons in the telencephalon [49]. All embryos and larvae were maintained at 28°C in E3 embryo medium (5 mM NaCl, 0.17 mM KCl, 0.33 mM MgSO4, 0.33 mM CaCl2), then E3 medium containing 0.003% 1-phenyl 2-thiourea (PTU) from approximately 8 hpf, and those media were refreshed daily. Embryos and larvae were removed from their chorions and used at the stage noted in each experiment. Fish were treated with 0.04% Tricine (MS-222, Ethyl m-Aminobenzoate Methanesulfonate, 14805-82, Nacalai Tesque) in the medium for anesthesia, chilled on ice to alleviate suffering, and then sacrificed by fixation in 4% Paraformaldehyde Phosphate Buffer Solution (09154-14, Nacalai Tesque) at 4 °C. All animal experiments for this study were approved by the Animal Experimentation Committee of Saitama Medical University (Approval No. 4250) and performed in accordance with the Regulation on Animal Experimentation at Saitama Medical University.

### Microinjection

Preparation and injection of antisense morpholino oligonucleotides (MOs) were carried out as previously described [31]. Briefly, MOs were diluted in 1× Danieau buffer (58 mM NaCl, 0.7 mM KCl, 0.4 mM MgSO4, 0.6 mM Ca(NO3)2, 5.0 mM HEPES, pH 7.6) and injected into 1 to 2-cell-stage embryos by using a microinjector FemtoJet 5247 (Eppendorf). The sequences and the concentrations of MOs (Gene Tools, OR, USA) used in this study are as follows: slc6a4a-AMO (splice-blocking antisense MO, MO1-slc6a4a, ZDB-MRPHLNO-141031-1, ZIRC) [69], 5’-ACGCACTTACATGCACTTACACATA -3’, at 4.178 ng/nl; slc6a4a-5mis (5mis; 5-bp-mismatched control), 5’- ACCCACTTAGATCCAGTTAGACATA -3’, at 4.198 ng/nl.

### Drug treatments

Preparation of paroxetine and 5-HT solution were carried out as described previously [47]. Briefly, paroxetine (Mw 365.83, P1977, Tokyo Chemical Industry (TCI), Tokyo, Japan), fluoxetine (Mw 345.79, F0750, TCI), fluvoxamine (Mw 434.41, F0858, TCI), sertraline (Mw 342.69, 193-16191, Fujifilm Wako Chemicals, Osaka, Japan) and 5-HT (Mw 212.68, S0370, TCI) were dissolved in dimethyl sulfoxide (DMSO) (Nacalai Tesque, 13407-45, Kyoto, Japan) at 10 mM for stock solutions and diluted with E3 medium to adjust the final concentration from 1–20 µM in 1% DMSO. Paroxetine pharmacokinetics is reported that the maximal blood concentration (C_max_) and the maximal time point (T_max_) are 1.93 ± 1.38 ng/mL and 4.61 ± 1.04 h, respectively, in adult healthy humans at a single dose of 10 mg paroxetine orally (Paxil Tablet, GSK). Our treatment with 1–20 µM paroxetine by bath application in E3 medium corresponds to 0.3748 and 7.4966 µg/mL *in vitro*, and thus approximately 200–3900 times higher than the C_max_ in the blood. Zebrafish embryos were dechorionated with forceps and immersed in the solution from 26-27 hpf, and then incubated at 28.5°C. Most of embryos treated under these conditions were survived (see Fig 2B), and those embryos were euthanized for further analyses or after social behavioral analysis.

### Immunohistochemistry

Whole-mount immunohistochemistry was carried out as previously described [31]. Briefly, embryos were fixed with 4% paraformaldehyde/PBS and incubated with primary antibodies at 4°C overnight, followed by incubation with secondary antibodies against rabbit or goat IgG conjugated with Alexa Flour 488 or 555 (Abcam, Cambridge, UK) together with DAPI (Thermo Fisher Scientific, MA, USA). The following primary antibodies were used in this study: anti-active Caspase-3 (rabbit, 1:500 dilution, 559565, BD Pharmingen, NJ, USA); anti-phospho histone H3 (pH3; 1:500 dilution, 9701, Cell Signaling, MA, USA); Sox2 antibody (rabbit, 1:500 dilution, GTX124477, GeneTex, Hsinchu, Taiwan); anti-Serotonin (goat, 1:500 dilution, ab66047, Abcam); SLC6A4/5-HTT (rabbit, 1:100 dilution, PA5-49572, Thermo Fisher Scientific); Cyclin D1 antibody [N1C3] (rabbit, 1:500 dilution, GTX108624, GeneTex). Following immunostaining, phosphate-buffered saline (PBS), the wash solution of embryos was gradually substituted by 30%, 50% and finally 70% glycerol/PBS for imaging under a microscope.

### RNA extraction and RT-qPCR

Total RNAs were extracted from whole embryos injected with MOs or treated with drugs using miRNeasy mini prep kit according to the manufacture’s protocols (Qiagen, Hilden, Germany). The total RNAs were used for cDNA synthesis using SuperScript IV reverse transcriptase (Thermo Fisher Scientific, MA, USA) with oligo dT primers, and obtained cDNAs were used for qPCR using Power SYBR Green Master Mix (Thermo Fisher Scientific, MA, USA) with gene specific primers. Primer sequences used in this study are: 5’-ATCTCTCCCGCTGAAGAGTTTTATG-3’ (slc6a4a-forward), 5’-GCCAGAGGTCTTGACCCCCTTC-3’ (slc6a4a-reverse), 5’-CGAGTCTAGTTCGAGTCCGC-3’ (sox2-forward), 5’-CGGGCAGGTACATACTGATC-3’ (sox2-rerverse), 5’-TCTGGAGGACTGTAAGAGGTATGC-3’ (rpl13α-forward) [68], 5’-AGACGCACAATCTTGAGAGCA-3’ (rpl13α-reverse) [68]. qPCR analysis was performed using PikoReal 96 Real-Time PCR system (Thermo Scientific, MA, USA). The expression levels of slc6a4a and sox2 relative to rpl13α were calculated by the 2^-ΔΔct^ method.

### Microscopy and imaging

Most of individuals were observed in lateral views under a microscopy unless otherwise noted. Each individual was mounted on a slide glass and adjusted for the left-light body axis to be parallel to the optical axis under a stereoscopic microscope as shown in S4 Fig. Whole body images were obtained using a stereoscopic microscope Leica MZ10F equipped with MC170 HD camera. Multicolor fluorescence images of the head and the tail were acquired using a confocal laser scanning microscope Zeiss LSM700 equipped with EC Plan-NEOFLUAR 20×/NA 0.5 and LD Plan-NEOFLUAR 40×/NA 0.6 objectives. The acquisition and the analysis of images were performed after validating our methods using different transgenic lines. To avoid the variance of fluorescence intensity by image-acquiring conditions, the conditions were initially determined using the control individuals per experiment, and all images were acquired under the same conditions. Each confocal image was obtained by compiling approximately 30–40 optical sections at 2.0 µm intervals per hemisphere using a sequential-scanning mode per each frame with 512 × 512 frame size and 8-bit depth. As a control, the mode of the fluorescence intensity frequency distribution within the ROI set in the hindbrain (see S4 Fig, B, C), which was 20, was regarded as the background level and removed from the compiled images. The same image processing was applied to all the images obtained by each experiment.

### Social behavioral analysis

A test arena (68 mm length × 39 mm width × 15 mm height) divided into three compartments (sibling, no sibling areas, 10-mm length on both sides; test area, 38-mm length in the center) was used to observe social behaviors of young fish (35–37 dpf). Swimming behaviors in the test arena were recorded by capturing movies using a cMOS camera (Blackfly S BFS-U3-32S4M, Point Grey) for 5 min at 50 frames per sec (fps).

Each experiment was performed in order of five steps as follows: (1) A fish was transferred to the test area and habituated for 5 min. (2) Without siblings, swimming behavior was recorded for 5 min (no sibling trial). (3) Siblings were transferred to the sibling area and habituated for 5min. (4) With siblings, swimming behavior was recorded for 5 min (sibling trial). (5) Captured movies were converted to avi format using ImageJ software, and tracking analysis of the swimming fish was performed using UMA Tracker software (Release 15). Social behaviors were analyzed by comparing (2) with (4) for Social Preference Index (SPI) calculated as follows: SPI = (Sibling frames – No sibling frames) / Total frames [50]. Sibling frames (blue) were defined as fish located in a half region on the side of sibling area, while no sibling frames (red) were defined as fish located in the other region in the test area. A value of SPI varied between –1 and 1 [50]. SPI difference is positive when SPI in the sibling trial is larger than the no sibling trial, and negative when SPI in the sibling trial is smaller than the no sibling trial. Difference in total distance was defined as the value in sibling trial subtracting that in no sibling trial.

### Quantification and statistical analysis

Body length, head area and Kaede, GFP or mCherry-positive areas and intensities were measured using Adobe Photoshop (2023, Ver. 24.4.1) and ImageJ (Ver. 1.53). Confocal stacks were converted into 2D images using sum intensity projection. To eliminate ectopic signals derived from noise, regions exceeding a defined threshold and comprising contiguous pixels with an area of 3 pixels or more were considered as foreground. The regions were defined as the fluorescence-positive area, and the area and fluorescence mean intensity were measured. An average of control values was calculated, and then each value of individuals was represented as a relative value to the average. Thus, the average of controls was denoted as 1.0 for every experiment. For statistical analysis of two or more samples, two-tailed, unpaired nonparametric Mann-Whitney U test or Kruskal-Wallis with Dunn’s multiple comparison tests were performed, unless otherwise noted, using GraphPad Prism 9 (Ver. 9.5.1). Permutation test was used to assess differences in SPI between sibling and non-sibling trials. For each comparison, individual SPI differences were randomly permuted between groups, and the test statistic was recalculated for each permutation. A total of 1000 permutations were performed to generate a null distribution. All results were calculated using SAS version 9.4 for Windows (SAS Institute, Cary, NC, USA).

## Acknowledgements

We are grateful to Hiroshi Handa, Keisuke Miyazawa, Masaki Hiramoto (Tokyo Medical University) for valuable discussions and generous support. We thank members of Anatomy and Experimental Animals (Saitama Medical University) and Hiromi Kazama (Tokyo Medical University) for help and technical support. Tg(*elavl3:Kaede*), Tg(*pou4fl-hsp70l:GFP*), Tg(*gfap:Gal4FF;UAS:GFP*) and SAGFF120A were provided from Zebrafish National BioResource Project. Tg(*pou4f3:GAL4;UAS:mCherry*) was kindly provided by Herwig Baier (Max-Planck Institute).

## Author contributions

T.S. conceived and designed the experiments. T.S., T.Ku. performed the anatomical and pharmacological experiments, and K.S., T.O. K.Y. performed behavioral experiments. T.S., K.S., S.T. analyzed the data. M.O., Y.S., Y.K. performed permutation test. T.S., S.T., T.Ka., and M.N. wrote and edited the manuscript.

## Supporting information

**S1 Fig. Distribution and knockdown of zebrafish SLC6A4a.** (**A**) Immunostaining with anti-SERT antibody for 30-hpf SLC6A4a-knockdown (KD) embryos by injection of slc6a4a antisense MO (AMO) and the control (5mis). (**B**) Quantification of immunofluorescent signals in SLC6A4a-KD embryos. (**C**) Expression of *slc6a4a* mRNAs in SLC6A4a-KD embryos by RT-qPCR. (**D**) Lateral views of 32-hpf embryos injected with antisense slc6a4a-AMO and the control slc6a4a-5mis. White line, head-tail body length. (**E**, **F**) Quantification of the body length (**E**) and the head area (**F**) of the injected embryos. (**G**) Lateral views of 32-hpf Tg(*elavl3:Kaede*) embryos injected with MOs (top) and those embryos immunostained for active Caspase-3 (bottom). (**H**) Quantification of the intensity of active Caspase-3 signals in the brain of the injected embryos. Scale bars: 200 µm (**D**); 100 µm (**A**, **G**). Data are presented as mean ± SEM. \**p* < 0.05, \*\*\**p* < 0.001, Mann-Whitney U test.

**S2 Fig. Development of morphology and the retinotectal projection in wild-type embryos.** (**A**) Lateral views of wild-type embryos at 50 hpf, 72 hpf, 5 dpf, and 7 dpf. White line, head-tail body length. (**B**, **C**) Quantification of the body length (**B**) and the head area (**C**) of wild-type embryos. (**D**) Lateral views of wild-type Tg(*pou4f1-hsp70l:GFP;pau4f3:GAL4;UAS:mCherry*) embryos at 72 hpf, 5 dpf, 7 dpf. GFP (green) expressed in tectal neurons and retinal axons. mCherry (red) expressed in retinal axons. (**E**, **F**) Quantification of GFP-positive (**E**) and mCherry-positive (**F**) areas in the optic tectum. (**G**, **H**) Quantification of the intensity of GFP (**G**) and mCherry (**H**) in the optic tectum. Scale bars: 200 µm (**A**); 100 µm (**D**). Data are presented as mean ± SEM. \**p* < 0.05, \*\**p* < 0.01, \*\*\**p* < 0.001, \*\*\*\**p* < 0.0001, ns, not significant. Kruskal-Wallis with Dunn’s multiple comparisons tests.

**S3 Fig. Effects of early paroxetine on embryogenesis and neurogenesis at 3 dpf.** (**A**) A schematic diagram of treatment with 1 µM paroxetine. (**B**) Lateral views of 73-hpf embryos treated with paroxetine and the control DMSO. (**C**, **D**) Quantification of the body length (**C**) and the head area (**D**) of the treated embryos. (**E**) A schematic diagram of treatment with 10 µM paroxetine. (**F**) Lateral views of 72-hpf embryos treated with paroxetine and the control DMSO. (**G**, **H**) Quantification of the body length (**G**) and the head area (**H**) of the treated embryos. (**I**) Lateral views of 72-hpf Tg(*pou4f1-hsp70l:GFP*) embryos treated with 10 µM paroxetine and the control DMSO. (**J**) Quantification of GFP intensity in the optic tectum of the treated embryos. White lines, head-tail body length. Parox, paroxetine. Scale bars: 200 µm (**B**, **F**); 100 µm (**I**). Data are presented as mean ± SEM. \**p* < 0.05, \*\**p* < 0.01, \*\*\**p* < 0.001, \*\*\*\**p* < 0.0001, Mann-Whitney U test.

**S4 Fig. Effects of early paroxetine on neurogenesis.** (**A**) Lateral views of a 50-hpf Tg(*neurod:EGFP*) embryo. Regions and lines for quantification of the intensity are indicated in the bottom image. (**B**) Intensity profiles of the yellow arrows in (**A**, bottom) in the optic tectum (green dots) and the hindbrain (gray dots). (**C**) Intensity frequencies of the regions (white squares in **A**, bottom) in the optic tectum (green dots) and the hindbrain (gray dots). (**D**) A schematic for the time course of treatment with paroxetine. (**E**) Lateral views of 50-hpf Tg(*elavl3:Kaede*) embryos treated with 1 µM, 10 µM paroxetine and the control DMSO. (**F**-**I**) Quantification of Kaede-positive area (**F**, **G**) and the intensity (**G**, **I**) in the optic tectum (**F**, **G**) and the telencephalon (**H**, **I**) of 10 µM paroxetine-treated embryos. (**J**) Expression of CyclinD1 proteins in the optic tectum. Images immunestained for CyclinD1 (red) merged with nuclear staining with DAPI (top) and binarized for CyclinD1 signals (bottom). (**K**) Lateral views of DMSO- and paroxetine-treated embryos immunostained for SOX2 (magenta). (**L**, **M**) Quantification of the expression of SOX2 proteins by immunohistochemistry (**L**), and *sox2* mRNA by RT-qPCR (**M**). Parox, paroxetine. Cb, cerebellum; e, eye; H, hindbrain; OT, optic tectum; T, telencephlon. Filled arrowhead, optic tectum; open arrowhead, telencephalon. Scale bars: 100 µm (**A**, **E**, **K**); 10 µm (**J**). Data are presented as mean ± SEM. \**p* < 0.05, \*\**p* < 0.01, Mann-Whitney U test.

**S5 Fig. Growth restriction and decreased neurogenesis by early transient paroxetine at 5 dpf.** (**A**) A schematic diagram of treatment with 10 µM paroxetine and 5-HT. (**B**) Lateral views of 5-dpf larvae treated with paroxetine, 5-HT and the control DMSO. White line, head-tail body length. (**C**, **D**) Quantification of the body length (**C**) and the head area (**D**) of the treated larvae. (**E**) Lateral views of 5-dpf Tg(*pou4f1-hsp70l:GFP*) larvae treated with 10 µM paroxetine and the control DMSO. (**F**, **G**) Quantification of GFP-positive area (**E**) and the intensity (**F**) in the optic tectum of the treated larvae. Parox, paroxetine. Scale bars: 200 µm (**B)**; 100 µm (**E**). Data are mean ± SEM. \**p* < 0.05, \*\*\**p* < 0.001, ns, not significant. Kruskal-Wallis with Dunn’s multiple comparisons tests (**C**, **D**), Mann-Whitney U test (**F**, **G**).

**S6 Fig. Decreased neurogenesis in the telencephalon and restored head morphology in paroxetine and 5-HT-treated 7-dpf larvae.** (**A**) A schematic diagram of treatment with 10 µM paroxetine and 5-HT. (**B**) Dorsal views of 48-hpf SAGFF120A;UAS:GFP embryos treated with paroxetine, 5-HT, and the control DMSO. (**C**) Quantification of the mean GFP intensities in the telencephalons of SAGFF120A;UAS:GFP embryos. (**D)** A schematic diagram of treatment with 1 µM paroxetine and 5-HT. (**E)** Lateral views of 7-dpf larvae treated with paroxetine, 5-HT and the control DMSO. White line, head-tail body length. (**F**, **G)** Quantification of the body length (**F**) and the head area (**G**) of the treated larvae. Parox, paroxetine. Scale bars: 100 µm (**B**); 200 µm (**E**). Data are mean ± SEM. \*\**p* < 0.01, \*\*\**p* < 0.001, ns, not significant. Kruskal-Wallis with Dunn’s multiple comparisons tests.

**S7 Fig. A summary of the results in this study.** The results obtained by treatment with paroxetine, other SSRIs and knockdown of SERT are shown in light blue, those by treatment with 5-HT are shown in magenta, and those with both SSRIs and 5-HT are shown in black on the background in pale blue.

**S1 Video. DMSO-treated fish in a no sibling trial.**

**S2 Video. DMSO-treated fish in a sibling trial.**

**S3 Video. Paroxetine-treated fish in a no sibling trial.**

**S4 Video. Paroxetine-treated fish in a sibling trial.**

**S5 Video. 5-HT-treated fish in a no sibling trial.**

**S6 Video. 5-HT-treated fish in a sibling trial.**

## References

1. Sestan N, State MW. Lost in Translation: Traversing the Complex Path from Genomics to Therapeutics in Autism Spectrum Disorder. Neuron. 2018;100(2):406–23. doi: 10.1016/j.neuron.2018.10.015.

2. Amaral DG, Schumann CM, Nordahl CW. Neuroanatomy of autism. Trends Neurosci. 2008;31(3):137–45. doi: 10.1016/j.tins.2007.12.005.

3. Iakoucheva LM, Muotri AR, Sebat J. Getting to the Cores of Autism. Cell. 2019;178(6):1287–98. doi: 10.1016/j.cell.2019.07.037.

4. Willsey HR, Willsey AJ, Wang B, State MW. Genomics, convergent neuroscience and progress in understanding autism spectrum disorder. Nat Rev Neurosci. 2022;23(6):323–41. doi: 10.1038/s41583-022-00576-7.

5. Al-Haddad BJS, Oler E, Armistead B, Elsayed NA, Weinberger DR, Bernier R, et al. The fetal origins of mental illness. Am J Obstet Gynecol. 2019;221(6):549–62. doi: 10.1016/j.ajog.2019.06.013.

6. Amgalan A, Andescavage N, Limperopoulos C. Prenatal origins of neuropsychiatric diseases. Acta Paediatr. 2021;110(6):1741–9. doi: 10.1111/apa.15766.

7. Wiggins LD, Rubenstein E, Daniels J, DiGuiseppi C, Yeargin-Allsopp M, Schieve LA, et al. A Phenotype of Childhood Autism Is Associated with Preexisting Maternal Anxiety and Depression. J Abnorm Child Psychol. 2019;47(4):731–40. doi: 10.1007/s10802-018-0469-8.

8. Howard LM, Molyneaux E, Dennis CL, Rochat T, Stein A, Milgrom J. Non-psychotic mental disorders in the perinatal period. Lancet. 2014;384(9956):1775–88. doi: 10.1016/S0140-6736(14)61276-9.

9. Hendrick V, Stowe ZN, Altshuler LL, Hwang S, Lee E, Haynes D. Placental passage of antidepressant medications. Am J Psychiatry. 2003;160(5):993–6. doi: 10.1176/appi.ajp.160.5.993.

10. Ewing G, Tatarchuk Y, Appleby D, Schwartz N, Kim D. Placental transfer of antidepressant medications: implications for postnatal adaptation syndrome. Clin Pharmacokinet. 2015;54(4):359–70. doi: 10.1007/s40262-014-0233-3.

11. Croen LA, Grether JK, Yoshida CK, Odouli R, Hendrick V. Antidepressant use during pregnancy and childhood autism spectrum disorders. Arch Gen Psychiatry. 2011;68(11):1104–12. doi: 10.1001/archgenpsychiatry.2011.73.

12. Velasquez JC, Goeden N, Bonnin A. Placental serotonin: implications for the developmental effects of SSRIs and maternal depression. Front Cell Neurosci. 2013;7:47. doi: 10.3389/fncel.2013.00047.

13. Millard SJ, Weston-Green K, Newell KA. The effects of maternal antidepressant use on offspring behaviour and brain development: Implications for risk of neurodevelopmental disorders. Neurosci Biobehav Rev. 2017;80:743–65. doi: 10.1016/j.neubiorev.2017.06.008.

14. Andalib S, Emamhadi MR, Yousefzadeh-Chabok S, Shakouri SK, Hoilund-Carlsen PF, Vafaee MS, et al. Maternal SSRI exposure increases the risk of autistic offspring: A meta-analysis and systematic review. Eur Psychiatry. 2017;45:161–6. doi: 10.1016/j.eurpsy.2017.06.001.

15. Nardozza LM, Caetano AC, Zamarian AC, Mazzola JB, Silva CP, Marcal VM, et al. Fetal growth restriction: current knowledge. Arch Gynecol Obstet. 2017;295(5):1061–77. doi: 10.1007/s00404-017-4341-9.

16. Dall’Asta A, Brunelli V, Prefumo F, Frusca T, Lees CC. Early onset fetal growth restriction. Matern Health Neonatol Perinatol. 2017;3:2. doi: 10.1186/s40748-016-0041-x.

17. Pels A, Knaven OC, Wijnberg-Williams BJ, Eijsermans MJC, Mulder-de Tollenaer SM, Aarnoudse-Moens CSH, et al. Neurodevelopmental outcomes at five years after early-onset fetal growth restriction: Analyses in a Dutch subgroup participating in a European management trial. Eur J Obstet Gynecol Reprod Biol. 2019;234:63–70. doi: 10.1016/j.ejogrb.2018.12.041.

18. Ranzil S, Ellery S, Walker DW, Vaillancourt C, Alfaidy N, Bonnin A, et al. Disrupted placental serotonin synthetic pathway and increased placental serotonin: Potential implications in the pathogenesis of human fetal growth restriction. Placenta. 2019;84:74–83. doi: 10.1016/j.placenta.2019.05.012.

19. Joseph RM, Korzeniewski SJ, Allred EN, O’Shea TM, Heeren T, Frazier JA, et al. Extremely low gestational age and very low birthweight for gestational age are risk factors for autism spectrum disorder in a large cohort study of 10-year-old children born at 23-27 weeks’ gestation. Am J Obstet Gynecol. 2017;216(3):304 e1- e16. doi: 10.1016/j.ajog.2016.11.1009.

20. Gaspar P, Cases O, Maroteaux L. The developmental role of serotonin: news from mouse molecular genetics. Nat Rev Neurosci. 2003;4(12):1002–12. doi: 10.1038/nrn1256.

21. Pourhamzeh M, Moravej FG, Arabi M, Shahriari E, Mehrabi S, Ward R, et al. The Roles of Serotonin in Neuropsychiatric Disorders. Cell Mol Neurobiol. 2022;42(6):1671–92. doi: 10.1007/s10571-021-01064-9.

22. Otte C, Gold SM, Penninx BW, Pariante CM, Etkin A, Fava M, et al. Major depressive disorder. Nat Rev Dis Primers. 2016;2:16065. doi: 10.1038/nrdp.2016.65.

23. Gross C, Zhuang X, Stark K, Ramboz S, Oosting R, Kirby L, et al. Serotonin1A receptor acts during development to establish normal anxiety-like behaviour in the adult. Nature. 2002;416(6879):396–400. doi: 10.1038/416396a.

24. Aimone JB, Li Y, Lee SW, Clemenson GD, Deng W, Gage FH. Regulation and function of adult neurogenesis: from genes to cognition. Physiol Rev. 2014;94(4):991–1026. doi: 10.1152/physrev.00004.2014.

25. Bonnin A, Goeden N, Chen K, Wilson ML, King J, Shih JC, et al. A transient placental source of serotonin for the fetal forebrain. Nature. 2011;472(7343):347–50. doi: 10.1038/nature09972.

26. Rosenfeld CS. The placenta-brain-axis. J Neurosci Res. 2021;99(1):271–83. doi: 10.1002/jnr.24603.

27. St-Pierre J, Laurent L, King S, Vaillancourt C. Effects of prenatal maternal stress on serotonin and fetal development. Placenta. 2016;48 Suppl 1:S66–S71. doi: 10.1016/j.placenta.2015.11.013.

28. Trowbridge S, Narboux-Neme N, Gaspar P. Genetic models of serotonin (5-HT) depletion: what do they tell us about the developmental role of 5-HT? Anat Rec (Hoboken). 2011;294(10):1615–23. doi: 10.1002/ar.21248.

29. Rea V, Van Raay TJ. Using Zebrafish to Model Autism Spectrum Disorder: A Comparison of ASD Risk Genes Between Zebrafish and Their Mammalian Counterparts. Front Mol Neurosci. 2020;13:575575. doi: 10.3389/fnmol.2020.575575.

30. Sakai C, Ijaz S, Hoffman EJ. Zebrafish Models of Neurodevelopmental Disorders: Past, Present, and Future. Front Mol Neurosci. 2018;11:294. doi: 10.3389/fnmol.2018.00294.

31. Sato T, Sato F, Kamezaki A, Sakaguchi K, Tanigome R, Kawakami K, et al. Neuregulin 1 Type II-ErbB Signaling Promotes Cell Divisions Generating Neurons from Neural Progenitor Cells in the Developing Zebrafish Brain. PLoS One. 2015;10(5):e0127360. doi: 10.1371/journal.pone.0127360.

32. Bourin M, Chue P, Guillon Y. Paroxetine: a review. CNS Drug Rev. 2001;7(1):25–47. doi: 10.1111/j.1527-3458.2001.tb00189.x.

33. Coleman JA, Green EM, Gouaux E. X-ray structures and mechanism of the human serotonin transporter. Nature. 2016;532(7599):334–9. doi: 10.1038/nature17629.

34. Coleman JA, Navratna V, Antermite D, Yang D, Bull JA, Gouaux E. Chemical and structural investigation of the paroxetine-human serotonin transporter complex. Elife. 2020;9. doi: 10.7554/eLife.56427.

35. Lillesaar C, Stigloher C, Tannhauser B, Wullimann MF, Bally-Cuif L. Axonal projections originating from raphe serotonergic neurons in the developing and adult zebrafish, Danio rerio, using transgenics to visualize raphe-specific pet1 expression. J Comp Neurol. 2009;512(2):158–82. doi: 10.1002/cne.21887.

36. Ito T, Handa H. Deciphering the mystery of thalidomide teratogenicity. Congenit Anom (Kyoto). 2012;52(1):1–7. doi: 10.1111/j.1741-4520.2011.00351.x.

37. Chen X, Liu H, Li Y, Zhang W, Zhou A, Xia W, et al. First-trimester fetal size, accelerated growth in utero, and child neurodevelopment in a cohort study. BMC Med. 2024;22(1):181. doi: 10.1186/s12916-024-03390-3.

38. Niell CM, Meyer MP, Smith SJ. In vivo imaging of synapse formation on a growing dendritic arbor. Nat Neurosci. 2004;7(3):254–60. doi: 10.1038/nn1191.

39. Hua JY, Smear MC, Baier H, Smith SJ. Regulation of axon growth in vivo by activity-based competition. Nature. 2005;434(7036):1022–6. doi: 10.1038/nature03409.

40. Sato T, Hamaoka T, Aizawa H, Hosoya T, Okamoto H. Genetic single-cell mosaic analysis implicates ephrinB2 reverse signaling in projections from the posterior tectum to the hindbrain in zebrafish. J Neurosci. 2007;27(20):5271–9. doi: 10.1523/JNEUROSCI.0883-07.2007.

41. Isa T, Marquez-Legorreta E, Grillner S, Scott EK. The tectum/superior colliculus as the vertebrate solution for spatial sensory integration and action. Curr Biol. 2021;31(11):R741–R62. Epub 2021/06/09. doi: 10.1016/j.cub.2021.04.001.

42. Xiao T, Baier H. Lamina-specific axonal projections in the zebrafish tectum require the type IV collagen Dragnet. Nat Neurosci. 2007;10(12):1529–37. Epub 2007/11/06. doi: 10.1038/nn2002.

43. Obholzer N, Wolfson S, Trapani JG, Mo W, Nechiporuk A, Busch-Nentwich E, et al. Vesicular glutamate transporter 3 is required for synaptic transmission in zebrafish hair cells. J Neurosci. 2008;28(9):2110–8. doi: 10.1523/JNEUROSCI.5230-07.2008.

44. Sato T, Takahoko M, Okamoto H. HuC:Kaede, a useful tool to label neural morphologies in networks in vivo. Genesis. 2006;44(3):136–42. doi: 10.1002/gene.20196.

45. Boldrini M, Hen R, Underwood MD, Rosoklija GB, Dwork AJ, Mann JJ, et al. Hippocampal angiogenesis and progenitor cell proliferation are increased with antidepressant use in major depression. Biol Psychiatry. 2012;72(7):562–71. doi: 10.1016/j.biopsych.2012.04.024.

46. Greig LC, Woodworth MB, Galazo MJ, Padmanabhan H, Macklis JD. Molecular logic of neocortical projection neuron specification, development and diversity. Nat Rev Neurosci. 2013;14(11):755–69. doi: 10.1038/nrn3586.

47. Ando H, Sato T, Ito T, Yamamoto J, Sakamoto S, Nitta N, et al. Cereblon Control of Zebrafish Brain Size by Regulation of Neural Stem Cell Proliferation. iScience. 2019;15:95–108. doi: 10.1016/j.isci.2019.04.007.

48. Scott EK, Mason L, Arrenberg AB, Ziv L, Gosse NJ, Xiao T, et al. Targeting neural circuitry in zebrafish using GAL4 enhancer trapping. Nat Methods. 2007;4(4):323–6. doi: 10.1038/nmeth1033.

49. Lal P, Tanabe H, Suster ML, Ailani D, Kotani Y, Muto A, et al. Identification of a neuronal population in the telencephalon essential for fear conditioning in zebrafish. BMC Biol. 2018;16(1):45. doi: 10.1186/s12915-018-0502-y.

50. Dreosti E, Lopes G, Kampff AR, Wilson SW. Development of social behavior in young zebrafish. Front Neural Circuits. 2015;9:39. doi: 10.3389/fncir.2015.00039.

51. Levkovitz Y, Gil-Ad I, Zeldich E, Dayag M, Weizman A. Differential induction of apoptosis by antidepressants in glioma and neuroblastoma cell lines: evidence for p-c-Jun, cytochrome c, and caspase-3 involvement. J Mol Neurosci. 2005;27(1):29–42. doi: 10.1385/JMN:27:1:029.

52. Schuster C, Fernbach N, Rix U, Superti-Furga G, Holy M, Freissmuth M, et al. Selective serotonin reuptake inhibitors--a new modality for the treatment of lymphoma/leukaemia? Biochem Pharmacol. 2007;74(9):1424–35. doi: 10.1016/j.bcp.2007.07.017.

53. Wei P, Jia H, Li R, Zhang C, Guo S, Wei S, et al. Fluvoxamine prompts the antitumor immune effect via inhibiting the PD-L1 expression on mice-burdened colon tumor. Cell Biol Int. 2023;47(2):439–50. doi: 10.1002/cbin.11936.

54. Hodge JM, Wang Y, Berk M, Collier FM, Fernandes TJ, Constable MJ, et al. Selective serotonin reuptake inhibitors inhibit human osteoclast and osteoblast formation and function. Biol Psychiatry. 2013;74(1):32–9. doi: 10.1016/j.biopsych.2012.11.003.

55. Reis G, Dos Santos Moreira-Silva EA, Silva DCM, Thabane L, Milagres AC, Ferreira TS, et al. Effect of early treatment with fluvoxamine on risk of emergency care and hospitalisation among patients with COVID-19: the TOGETHER randomised, platform clinical trial. Lancet Glob Health. 2022;10(1):e42–e51. doi: 10.1016/S2214-109X(21)00448-4.

56. Hashimoto Y, Suzuki T, Hashimoto K. Mechanisms of action of fluvoxamine for COVID-19: a historical review. Mol Psychiatry. 2022;27(4):1898–907. doi: 10.1038/s41380-021-01432-3.

57. Asatsuma-Okumura T, Ando H, De Simone M, Yamamoto J, Sato T, Shimizu N, et al. p63 is a cereblon substrate involved in thalidomide teratogenicity. Nat Chem Biol. 2019;15(11):1077–84. doi: 10.1038/s41589-019-0366-7.

58. Miller MT, Stromland K, Ventura L, Johansson M, Bandim JM, Gillberg C. Autism with ophthalmologic malformations: the plot thickens. Trans Am Ophthalmol Soc. 2004;102:107–20; discussion 20-1.

59. Sato T, Ito T, Handa H. Cereblon-Based Small-Molecule Compounds to Control Neural Stem Cell Proliferation in Regenerative Medicine. Front Cell Dev Biol. 2021;9:629326. doi: 10.3389/fcell.2021.629326.

60. Ito T, Yamaguchi Y, Handa H. Exploiting ubiquitin ligase cereblon as a target for small-molecule compounds in medicine and chemical biology. Cell Chem Biol. 2021;28(7):987–99. doi: 10.1016/j.chembiol.2021.04.012.

61. Kriegstein A, Alvarez-Buylla A. The glial nature of embryonic and adult neural stem cells. Annu Rev Neurosci. 2009;32:149–84. doi: 10.1146/annurev.neuro.051508.135600.

62. Suh H, Consiglio A, Ray J, Sawai T, D’Amour KA, Gage FH. In vivo fate analysis reveals the multipotent and self-renewal capacities of Sox2+ neural stem cells in the adult hippocampus. Cell Stem Cell. 2007;1(5):515–28. doi: 10.1016/j.stem.2007.09.002.

63. Sahay A, Hen R. Adult hippocampal neurogenesis in depression. Nat Neurosci. 2007;10(9):1110–5. doi: 10.1038/nn1969.

64. Persico AM, Baldi A, Dell’Acqua ML, Moessner R, Murphy DL, Lesch KP, et al. Reduced programmed cell death in brains of serotonin transporter knockout mice. Neuroreport. 2003;14(3):341–4. doi: 10.1097/00001756-200303030-00009.

65. Salichon N, Gaspar P, Upton AL, Picaud S, Hanoun N, Hamon M, et al. Excessive activation of serotonin (5-HT) 1B receptors disrupts the formation of sensory maps in monoamine oxidase a and 5-ht transporter knock-out mice. J Neurosci. 2001;21(3):884–96. doi: 10.1523/JNEUROSCI.21-03-00884.2001.

66. Upton AL, Ravary A, Salichon N, Moessner R, Lesch KP, Hen R, et al. Lack of 5-HT(1B) receptor and of serotonin transporter have different effects on the segregation of retinal axons in the lateral geniculate nucleus compared to the superior colliculus. Neuroscience. 2002;111(3):597–610. doi: 10.1016/s0306-4522(01)00602-9.

67. Avino TA, Barger N, Vargas MV, Carlson EL, Amaral DG, Bauman MD, et al. Neuron numbers increase in the human amygdala from birth to adulthood, but not in autism. Proc Natl Acad Sci U S A. 2018;115(14):3710–5. doi: 10.1073/pnas.1801912115.

68. Westerfield M. THE ZEBRAFISH BOOK, 5th ed. A guide for the laboratory use of zebrafish (Danio rerio). Eugene: University of Oregon Press; 2007.

69. Waugh TA, Horstick E, Hur J, Jackson SW, Davidson AE, Li X, et al. Fluoxetine prevents dystrophic changes in a zebrafish model of Duchenne muscular dystrophy. Hum Mol Genet. 2014;23(17):4651–62. doi: 10.1093/hmg/ddu185.

70. Rassier GT, Silveira TLR, Remiao MH, Daneluz LO, Martins AWS, Dellagostin EN, et al. Evaluation of qPCR reference genes in GH-overexpressing transgenic zebrafish (Danio rerio). Sci Rep. 2020;10(1):12692. doi: 10.1038/s41598-020-69423-y.

71. Lukaszewicz AI, Anderson DJ. Cyclin D1 promotes neurogenesis in the developing spinal cord in a cell cycle-independent manner. Proc Natl Acad Sci U S A. 2011;108(28):11632–7. doi: 10.1073/pnas.1106230108.

72. Schaefer T, Steiner R, Lengerke C. SOX2 and p53 Expression Control Converges in PI3K/AKT Signaling with Versatile Implications for Stemness and Cancer. Int J Mol Sci. 2020;21(14). doi: 10.3390/ijms21144902.

73. Chen R, Zou J, Liu J, Kang R, Tang D. DAMPs in the immunogenicity of cell death. Mol Cell. 2025;85(20):3874–89. doi: 10.1016/j.molcel.2025.09.007.

74. Gahtan E, Tanger P, Baier H. Visual prey capture in larval zebrafish is controlled by identified reticulospinal neurons downstream of the tectum. J Neurosci. 2005;25(40):9294–303. doi: 10.1523/JNEUROSCI.2678-05.2005.

