## Supplementary figures and images for "Paroxetine-induced transient apoptosis with delayed neurogenesis induces brain remodeling in developing zebrafish"

### Supplemental Figure S1

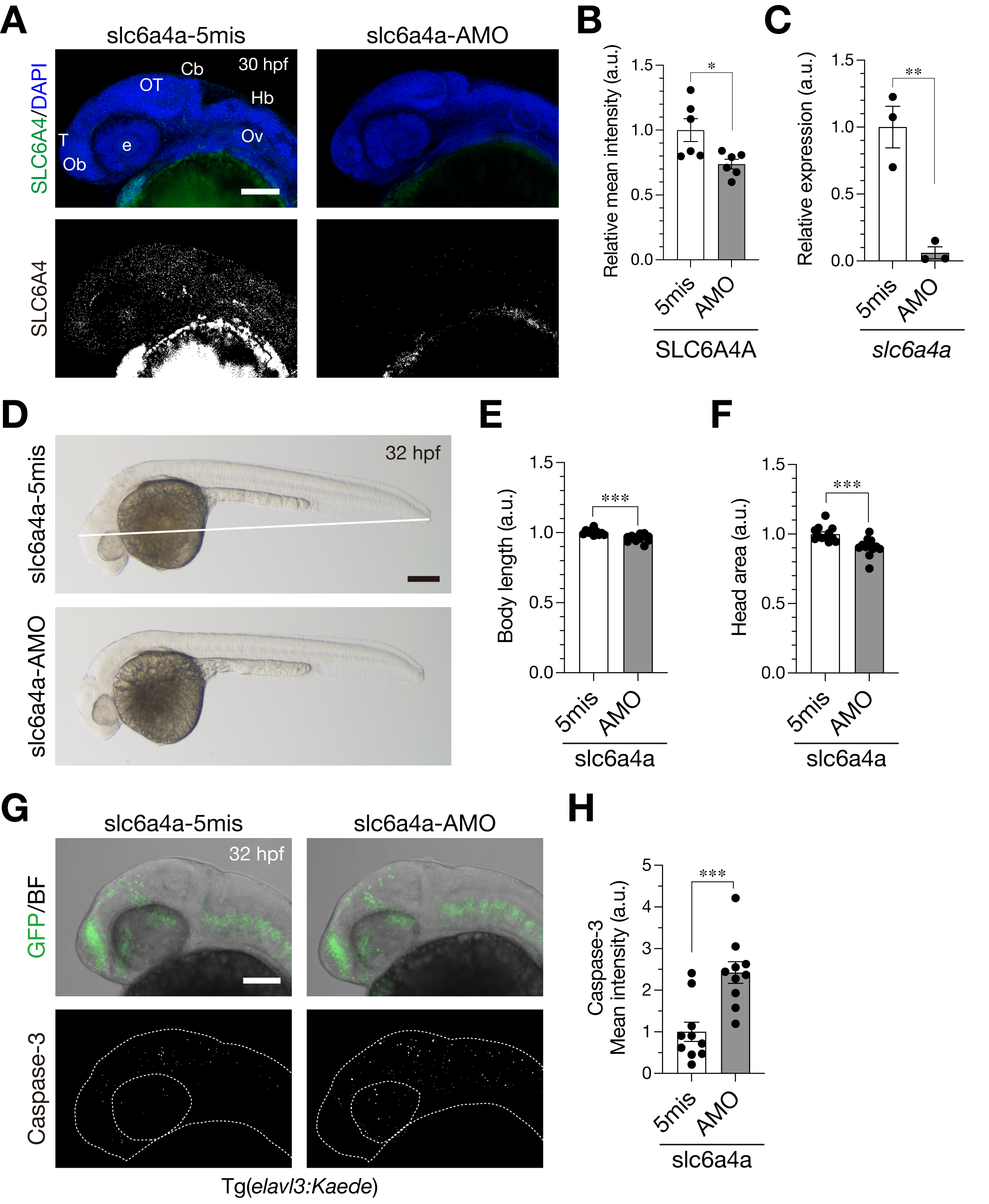

### Supplemental Figure S2

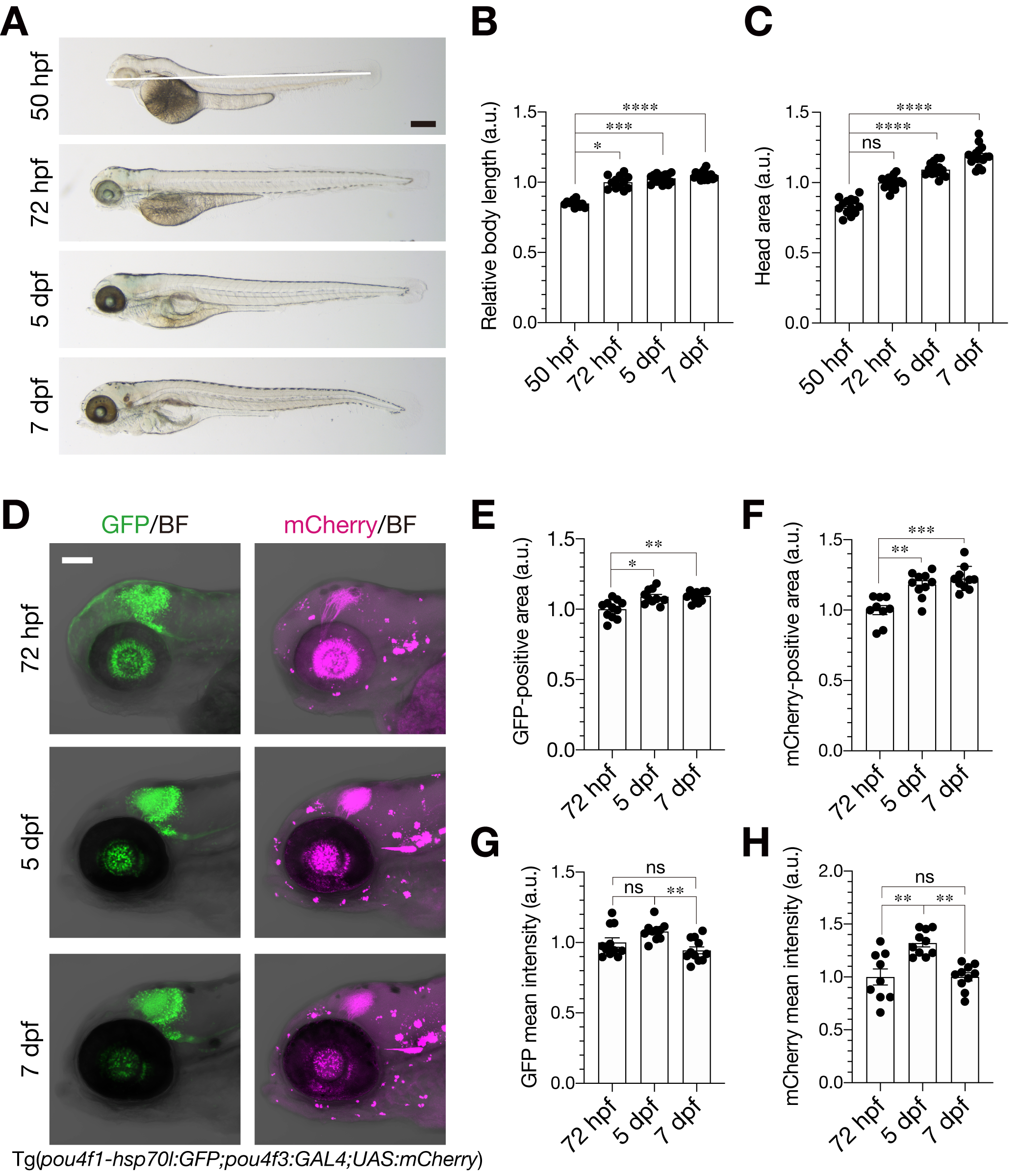

### Supplemental Figure S3

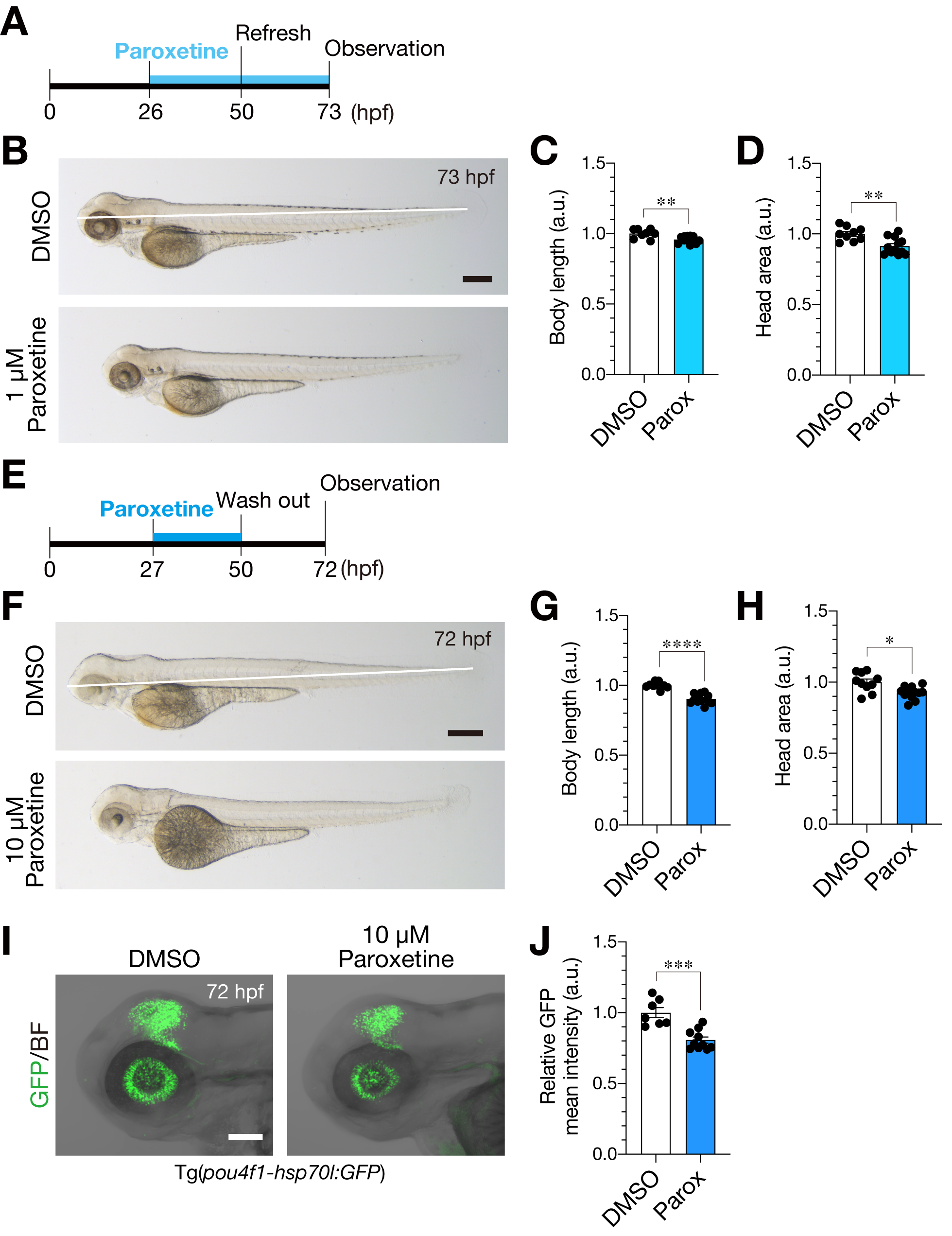

### Supplemental Figure S4

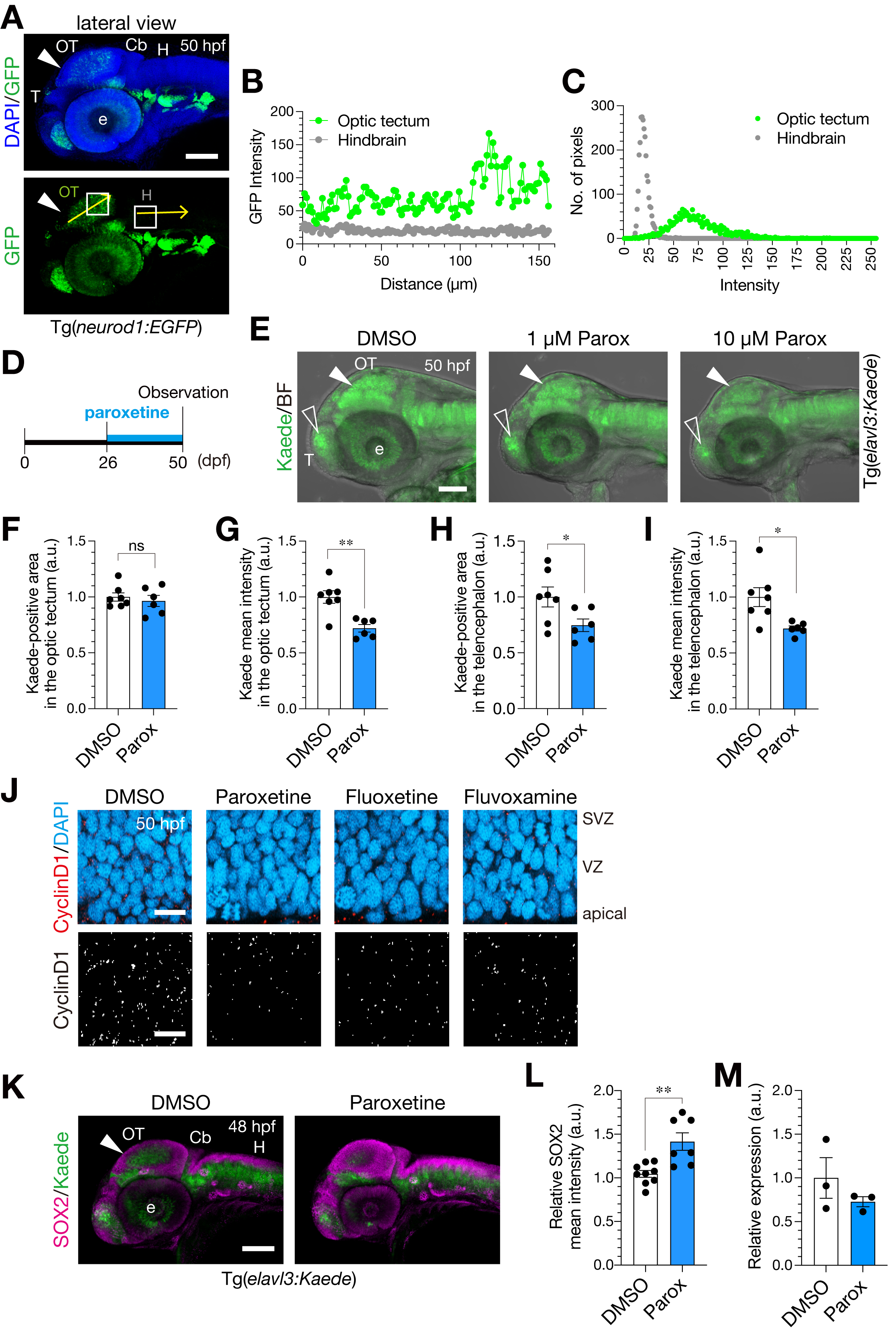

### Supplemental Figure S5

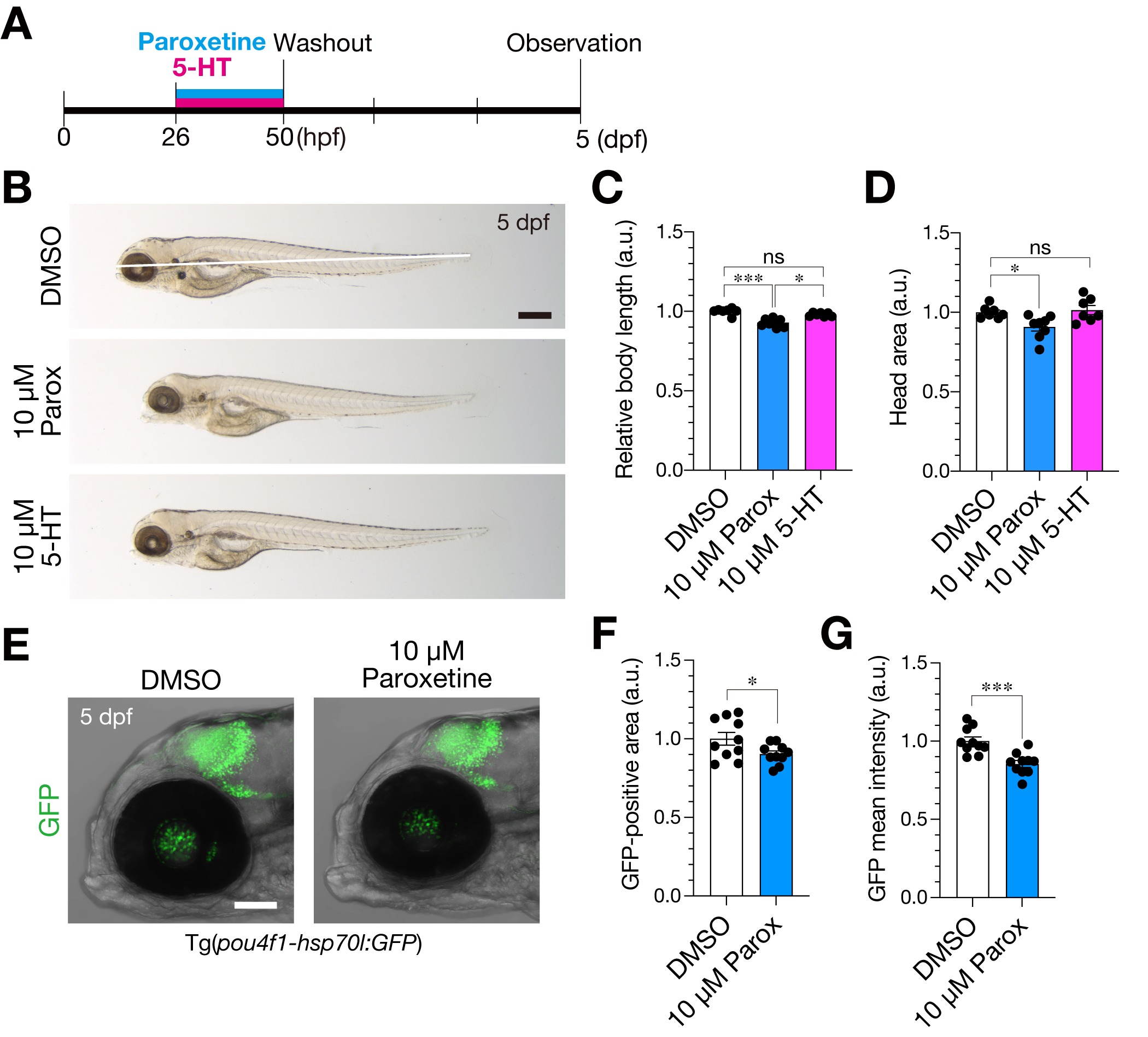

### Supplemental Figure S6

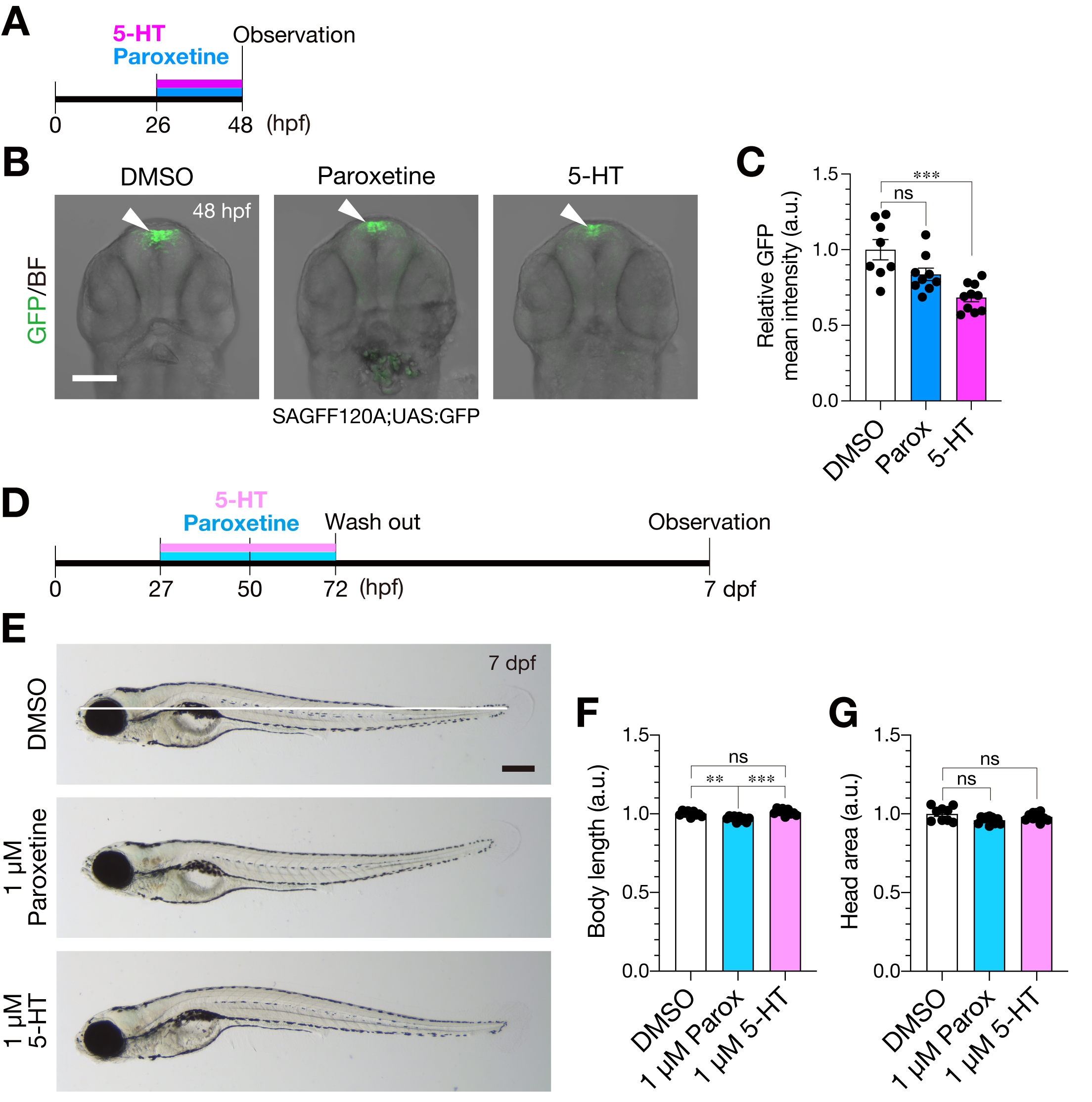

### Supplemental Figure S7

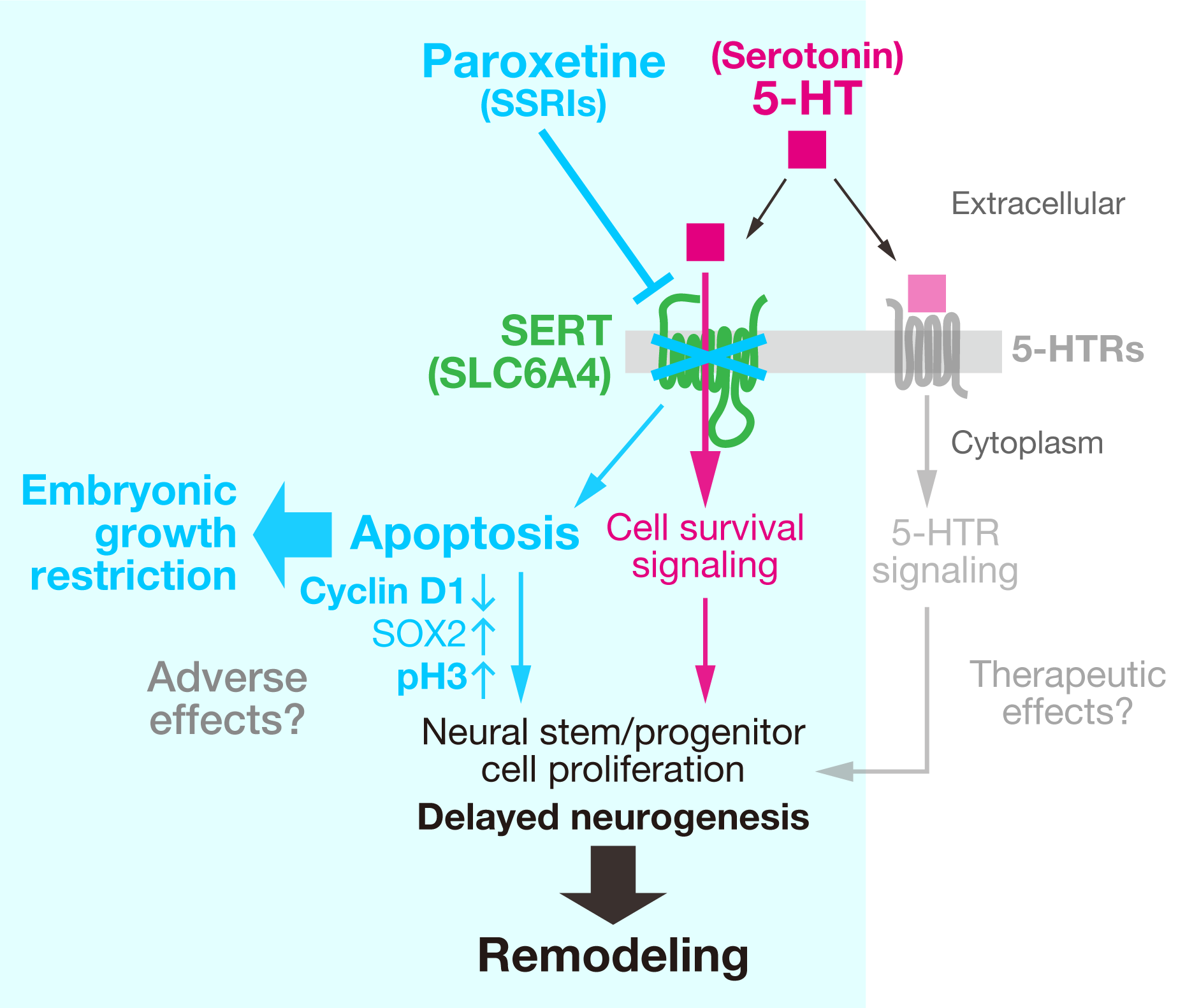
